# SpottedPy quantifies relationships between spatial transcriptomic hotspots and uncovers new environmental cues of epithelial-mesesenchymal plasticity in breast cancer

**DOI:** 10.1101/2023.12.20.572627

**Authors:** Eloise Withnell, Maria Secrier

## Abstract

Spatial transcriptomics is revolutionising the exploration of intratissue heterogeneity in cancer, yet capturing cellular niches and their spatial relationships remains challenging. We introduce SpottedPy, a Python package designed to identify tumour hotspots and map spatial interactions within the cancer ecosystem. Using SpottedPy, we examine epithelial-mesenchymal plasticity in breast cancer and highlight stable niches associated with angiogenic and hypoxic regions, shielded by CAFs and macrophages. Hybrid and mesenchymal hotspot distribution followed transformation gradients reflecting progressive immunosuppression. Our method offers flexibility to explore spatial relationships at different scales, from immediate neighbours to broader tissue modules, providing new insights into tumour microenvironment dynamics.

## BACKGROUND

The spatial organisation of cancer cells and their interactions with immune and stromal cells in their environment are instrumental in dictating tumour behaviour and progression(1). Recent developments in sequencing technologies that are able to profile large areas of the entire tissue in minute detail, such as spatial transcriptomics(2,3), are increasingly enabling us to gain a more comprehensive understanding of the complex ecosystem of tumour microenvironments (TME)(4),(5). Profiling the cancer tissue spatially allows us to explore the tumour architecture and its heterogeneity in detail, paving the way to deciphering the crosstalk between tumour cells and their surroundings and opening up new therapeutic opportunities(6),(7).

Several studies to date have demonstrated the advantages of spatial transcriptomics in delineating major tissue domains with distinct cell composition(8), cancer hallmarks(9), immunosuppressive hubs(10,11), entire tumour ecotypes with divergent clinical outcomes(12) or the impact of specific drugs on inhibiting tumour progression(13). However, focusing on areas of the tissue that are relevant for a specific biological question and surveying relationships between cell populations at the right scale remains a challenge in these datasets. To delineate biologically meaningful tissue areas using spatial transcriptomics, some of the current analytical methods, such as SpaGCN(14) and BayesSpace(15) focus on unsupervised clustering of gene expression. Approaches like NeST(16) or GASTON(17) take this one step further and incorporate a nested structure or topography metrics to outline hierarchically organised co-expression hotspots aligning with tissue histology. Given that similar cells often cluster together(18),(19), methods that can reliably detect statistically significant spatial clusters are important in reinforcing the accuracy of cell states determined from continuous signatures. CellCharter(20) capitalises on this concept and uses Gaussian mixture models to identify stable clusters representing spatial niches exhibiting distinct shapes and cell plasticity. More targeted clustering approaches employing user-defined signatures or cell types have recently been implemented in Voyager(21) and Monkeybread(22). Such methods can play a significant role in enhancing the interpretation of cell types that are inferred through spatial transcriptomic deconvolution techniques, but flexibly exploring spatial units at different scales remains difficult.

When it comes to assessing the spatial proximity of different clusters, methods that infer this through co-enrichment within the immediate neighbourhood have been implemented in packages like Squidpy(23). However, there is a lack of methods that calculate a differential spatial relationship between cell types or signatures of interest, e.g. hypoxia. Additionally, current approaches lack analytical methods to define and compare shorter and longer-range interactions between specific areas or cell populations of interest. Given that the scale of certain biological processes in cancer, such as hypoxia, remains elusive, relying solely on conventional spot neighbourhood-centric methods might obscure complex, spatial interactions. This is known as the Modifiable Areal Unit Problem (MAUP) in geostatistics(24), where spatial data patterns are observed to shift contingent on the size and shape of the spatial analysis units. Whilst a growing number of methods address the need for multi-scale analysis (11,23–26), the effect of changing spatial units has generally been underexplored in spatial biology(18).

Here, we build on the use of key ideas in the geostatistics field within spatial biology as previously demonstrated by Voyager(21) to devise an analytical method tailored to interrogate spatial relationships at various scales within 10X Visium transcriptomic datasets. Our approach defines areas densely inhabited by particular cell types or marked by user-defined gene signatures (hotspots) and areas depleted of cell types or gene signatures (coldspots), and statistically evaluates the proximity of such areas to other predefined hotspots or coldspots. We additionally compare the spatial relationships detected using the hotspot approach to those observed when looking only at the immediate neighbourhood of individual spots. We assess how the relationships between these variables change when varying the hotspot size or the neighbourhood size surrounding a spot. Importantly, we allow users to perform differential spatial analysis between two signatures or cell types of interest. For example, we can answer questions such as “which immune hotspots are significantly closer to mesenchymal hotspots compared to epithelial hotspots?” We have implemented this method in the Python package SpottedPy, available at https://github.com/secrierlab/SpottedPy.

To highlight the potential of our method, we focus on a key process underlying cancer progression, the epithelial-to-mesenchymal transition (EMT). During EMT, polarised epithelial cells undergo multiple molecular changes and lose their identity to acquire a mesenchymal phenotype(27). The interplay between EMT and the tumour microenvironment (TME) is multifaceted: while the TME is believed to be an inducer of EMT, mesenchymal tumour cells potentially influence the TME(28),(29). This dynamic is further complicated by the nature of EMT, which is not merely a dichotomous event. Current research suggests that EMT is a spectrum, varying from a continuous gradient to distinct, discrete stages(30),(31). Depending on the context, cells undergoing this transition can be locked in an EMT state, or alternate between a large landscape of EMT states, a phenomenon called epithelial-mesenchymal plasticity (EMP)(32). We have previously shown that the tumour spatial organisation follows EMP gradients and that hybrid and mesenchymal cells establish distinct interactions with the TME in small datasets of spatially-profiled breast tumours(33). However, the spatial organisation and interactions of cells across this EMP spectrum within the tumour milieu remain largely undefined at a larger scale and could offer great potential in developing therapies that exploit cell intrinsic or microenvironmental vulnerabilities linked with this process. Here, we showcase the capability of SpottedPy to unveil new relationships between EMT hotspots and the TME in breast cancer spatial transcriptomics data. By facilitating multi-scale comparisons of tumour and TME relationships, we yield rigorous evidence of spatial dynamics and offer an interpretable and intuitive measure of interactions between tumour cells and their environment at flexible scales.

## RESULTS

### SpottedPy: a tool to investigate biological modules and spatial relationships at different scales

From direct cell-cell interactions to immediate neighbourhoods and even across larger modules, cancer cell evolution is impacted at different scales by its environment (**Fig. 1a**). However, exploring this landscape flexibly and determining the areas within the tissue where these effects are most prominent is not straightforward. Whilst neighbourhood enrichment is commonly used in the field(23),(25),(28), analysing continuous expression signatures and examining of how the size of the neighbourhood influences spatial relationships are areas that remain underdeveloped. Furthermore, inspecting the immediate neighbourhood of cells of interest versus broader hotspots within the tissue will yield different insights into the tissue architecture and organisation, and this warrants further investigation.

**Figure 1:**
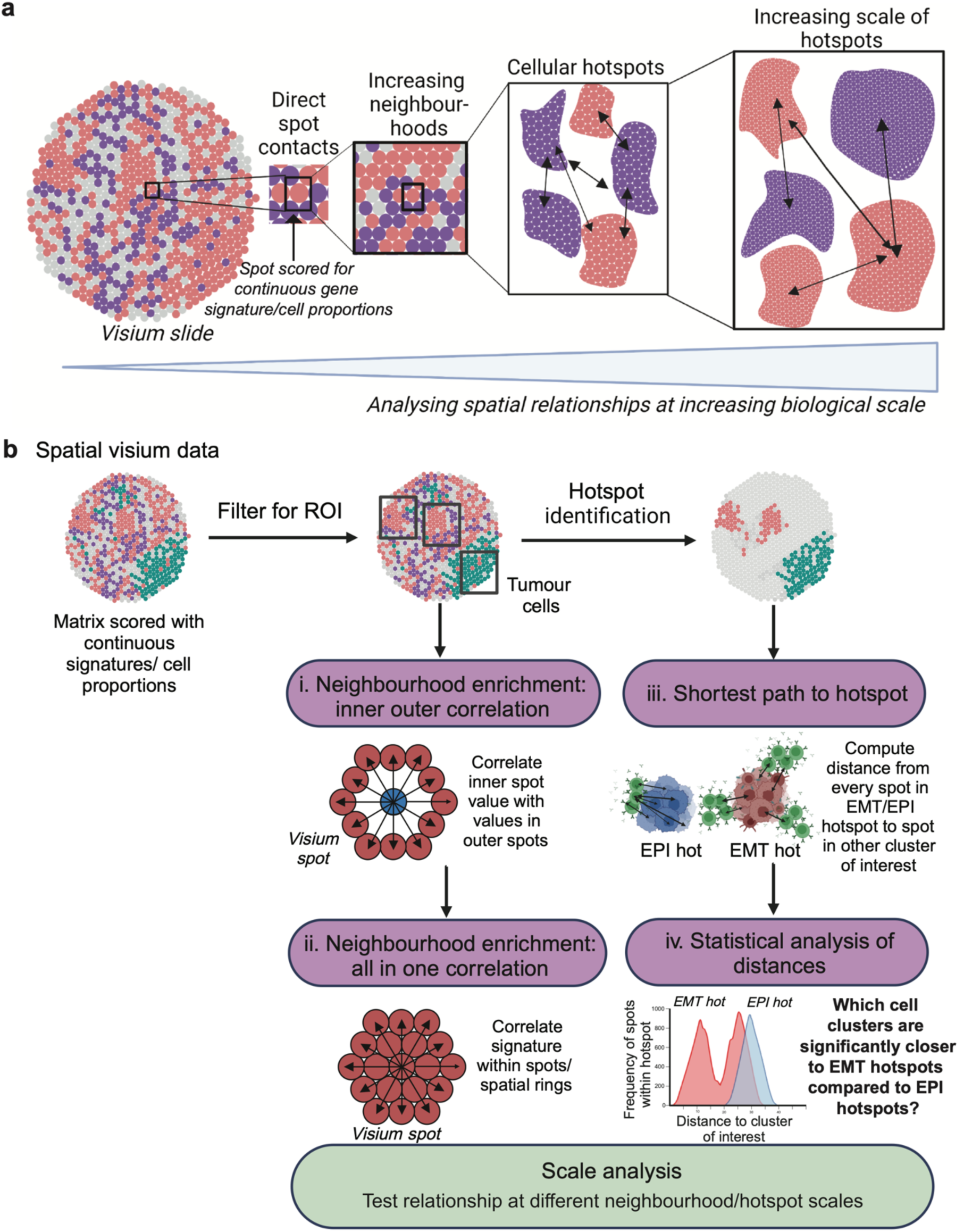
SpottedPy provides a multi-scale approach to analyse spatial transcriptomic relationships. (**a**) Overview of spatial scales captured in the SpottedPy workflow, from direct cellular contacts to broader cellular hotspots. Figure created with BioRender.com. (**b**) SpottedPy workflow overview. Visium spatial transcriptomic data is loaded as a pre-processed *AnnData* object where there is the option to select the region of interest (ROI) within the slide e.g., *AnnData.obs* column labelled with tumour cells. The default spatial analytics include: (i) Neighbourhood enrichment: inner outer correlation, which correlates cell prevalence or signatures in individual spots with their immediate neighbourhood, (ii) Neighbourhood enrichment: all in one correlation, which correlates cell prevalence of signatures within a spot or spatial unit (iii) Shortest path to hotspot, which calculates the minimum distance between each spot within a selected hotspot and the nearest spot in other hotspots, (iv) Statistical analysis of distances, which compares distances from a reference hotspot to another hotspot of interest, and assesses the statistical significance of the relationships. Scale analysis allows us to compare relationships defined at different scales in both approaches, either by increasing the number of rings included for neighbourhood enrichment or increasing the hotspot size. The outputs for the different modules include various plots to highlight the relationships. Figure created with BioRender.com.

We introduce SpottedPy, a Python package that enables a statistically principled interrogation of biological modules and cell-cell relationships at different scales in spatially profiled tissue (**Fig. 1b**).

This method encompasses:

- **Neighbourhood enrichment analysis:** Our method introduces a function designed to examine correlations between cell states, populations or processes within individual spatial transcriptomics spots and their immediate neighbourhood (**Fig. 1bi-ii**). In this context, a ‘neighbourhood’ is delineated as a ring encompassing six Visium spots around a designated central spot, which we calculate by treating the spatial spots as a network. We offer functionality to test how a signature effects its direct neighbourhood (inner-outer correlation), or within neighbourhood units (all-in-one correlation).
- **Hotspot identification:** We have implemented the Getis-Ord G* statistic to identify spatial clusters of continuous gene signatures across the tumour slide (**Fig. 1b).** Users can flexibly filter for specific regions within the slide to focus on when creating hotspots. By comparing defined regions with high or low expression or cell type signatures against a null hypothesis of random distribution, this analysis reveals statistically significant ‘hotspots’ or ‘coldspots’. Hotspots represent areas with a high concentration of a particular cell type or signature, suggesting an aggregation that is unlikely to be due to chance. Conversely, coldspots indicate regions where the target cells or signatures are scarce, also beyond what would be expected in a random distribution. We provide functionality to test enrichment of specific signatures within hotspots and coldspots.
- **Distance statistics:** SpottedPy includes analytical tools to measure and interpret the distances between identified clusters, such as between tumour and immune hotspots. The primary method calculates the *shortest path to a hotspot*, defined as the minimum distance from any point within a defined hotspot to the nearest point within a specified comparison hotspot (**Fig. 1biii**). Importantly, SpottedPy allows the user to compare distance distributions to key hotspots, for example, finding the hotspots that are significantly closer to mesenchymal hotspots than epithelial hotspots (or other areas that can be considered as a reference) (**Fig. 1biv).** Importantly, SpottedPy assigns a statistical significance to these proximity measures to determine if observed patterns are likely to occur just by chance. To statistically analyse the relationships across multiple slides, we use generalised estimating equations. SpottedPy allows the user to test either the minimum, mean or median distance from each hotspot, or assess all distances from each spot within a hotspot.
- **Scale/sensitivity analysis:** We provide the ability to systematically evaluate how cell-cell relationships evolve within the tissue as we vary the size of the neighbourhood or of the hotspot of interest. For the neighbourhood enrichment approach, this can be assessed by varying the number of concentric rings around the central spot. For the hotspot approach, SpottedPy recalculates the Getis-Ord G* statistic with varying neighbourhood sizes, enabling the identification of clusters at different spatial scales. By analysing how the distances between hotspots change with neighbourhood size, the package can illuminate shifting spatial relationships, providing insights into how biological entities interact across different scales. Additionally, SpottedPy allows users to examine how cluster relationships change when modifying the significance threshold for identifying hotspots with the Getis-Ord G* statistic.

To highlight the potential of our method, we employ SpottedPy to investigate the relationships between tumour cells undergoing EMT and the TME in 12 breast cancer slides profiled using the 10x Genomics Visium spatial transcriptomics platform, integrated from Wu et al^18^, Barkley et al^24^ and the 10x Genomics website^25^. To infer individual cell types within the slides, we performed cell deconvolution using the Cell2location method(35) and a scRNA-seq reference of annotated breast cancer cell population profiles from 123,561 cells^13^. We scored the tumour cells in the scRNA-seq dataset with a defined epithelial (EPI) and an epithelial-to-mesenchymal transition (EMT) signature (see Methods), and used Gaussian mixture modelling to assign a state to the tumour cells (36,37). To more precisely identify tumour cells within the spatial transcriptomic data, which tend to show vast expression variability, we employed the copy number inference tool STARCH(38) and only kept spots that showed evidence of copy number changes, which are likely to be tumour-specific. We validated the STARCH results by comparing to publicly available pathologist annotated slides (39,40) (**Supplementary Fig. 1a-b**). Furthermore, to explore the heterogeneity of EMT stable states established during the development and progression of breast cancer, we employed the discrete EMT states recently defined by Brown et al(41), encompassing an epithelial phenotype, two intermediate (hybrid) states (EM2 and EM3), a late intermediate quasi-mesenchymal state (M1) and a fully mesenchymal state (M2).

### The spatial landscape of EMT and associated tumour hallmarks

Firstly, we aimed to explore the cellular environment that is conducive to E/M progression. To do so, we focused solely on tumour hotspots and two key cellular hallmarks that have been previously linked with EMT, hypoxia and angiogenesis. Hypoxia, characterised by low oxygen levels, has been long recognized as a key enabler of tumorigenic processes(42). Under hypoxic conditions, tumour cells stabilise hypoxia-inducible factors (HIFs), primarily HIF-1α, which promotes angiogenesis(43), the formation of new blood vessels from pre-existing vasculature, to re-establish oxygen supply. Hypoxia has been shown to induce EMT and resistance to therapy(44), and therefore understanding how such relationships develop spatially within the tissue can help devise localised therapies that can interrupt these interactions in breast cancer.

Within individual spatial transcriptomics slides, we used SpottedPy to delineate tumour areas, and further identified EMT hotspots within these areas using using the EMT state as assigned using Cell2location (**Fig. 2a).** A certain degree of heterogeneity in the number and spatial distribution of EMT hotspots could be observed across the cohort (**Supplementary Fig. 1c).** Notably, Slides 3, 7 and 10 exhibited the greatest dispersion, which was independent of cancer subtype (TNBC and ER+ HER2+). This dispersion did not influence overall EMT relationships with other cell types (**Supplementary Fig. 2a**). In contrast, Slides 1 and 5 showed minimal dispersion, also without affecting EMT relationships with other cell types. The distributions of individual EMT and EPI signature scores per slide are highlighted in **Supplementary Fig. 1c-d**. The EPI signature typically followed a normal distribution across spatial tumour spots, whereas the EMT signature exhibited a positive skew, in line with this state being expected to be rarer within the primary tumours. Slides 0, 1, and 9 display more pronounced skewness for the EPI signature, suggesting a more advanced stage of EMT transformation within these tumours, which was not an effect of the cancer subtype as these slides present different breast cancer pathologies.

**Figure 2:**
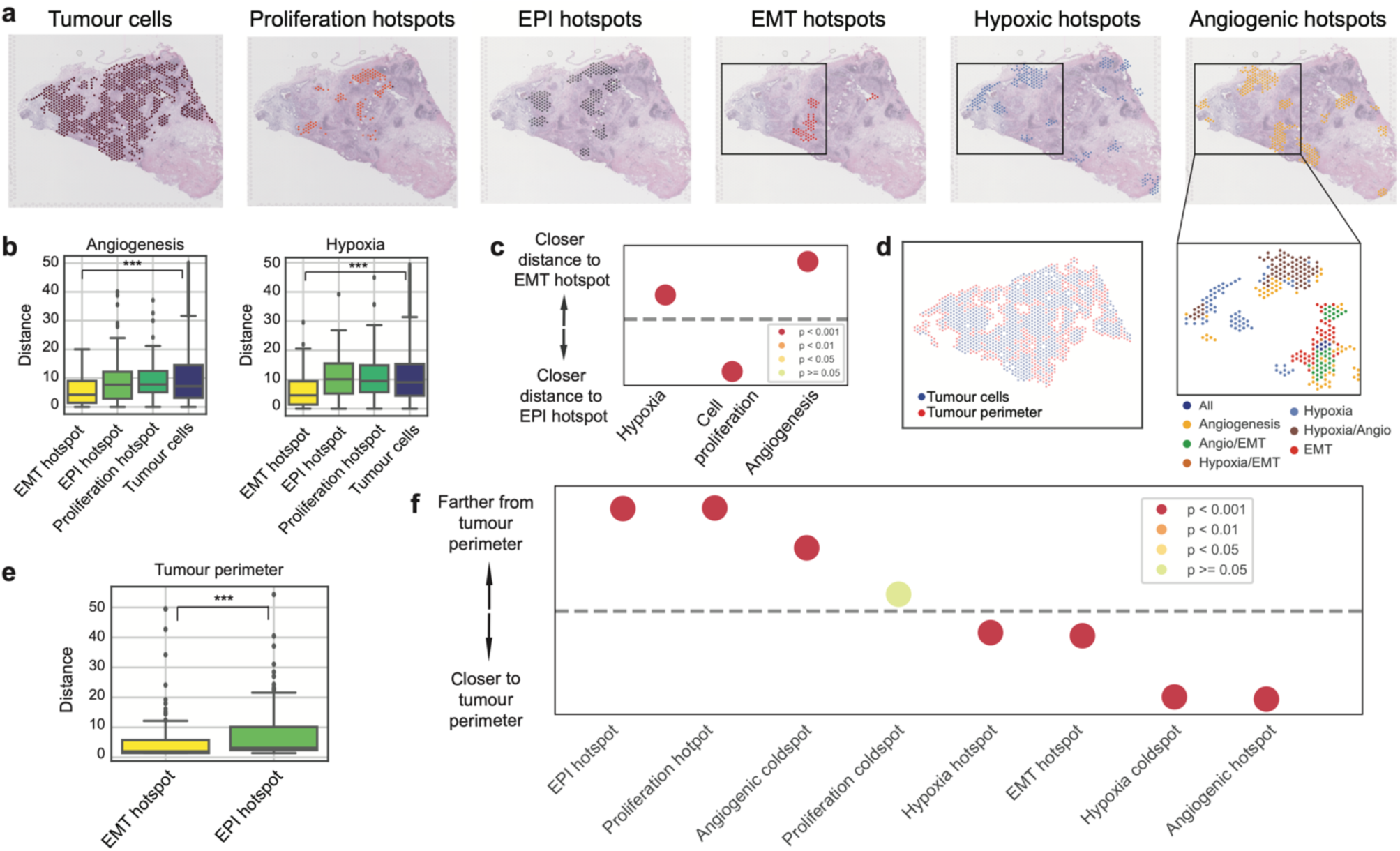
The spatial interplay between EMT progression and cancer hallmarks. (**a**) Spatial transcriptomics slide (Slide 0) highlighting from left to right: tumour spots, proliferation hotspots, EPI hotspots, EMT hotspots, hypoxic hotspots and angiogenic hotspots identified by SpottedPy. The black square indicates a representative area where the close proximity of EMT, angiogenic and hypoxic hotspots is depicted. (**b**) Distances from angiogenic (left) and hypoxic (right) hotspots to EMT hotspots, EPI hotspots, proliferative hotspots and the average tumour cell, respectively, averaged across all 12 samples (*** p<0.001). (**c**) Differences in proximity between EMT hotspots/EPI hotspots and hypoxic, proliferative and angiogenic regions, summarised across the 12 slides. The dashed line represents no difference in relative distance to EMT hotspots or EPI hotspots. The dots situated above the dashed line indicate hallmarks that are significantly closer to EMT hotspots. The colours indicate the p-value ranges obtained from the Student’s t test for differences in distance to EMT hot/cold areas. (**d**) Spatial plot depicting the tumour perimeter in red, tumour cells in blue and the microenvironment in grey. (**e**) Distance from the tumour perimeter to EMT hotspots and EPI hotspots, respectively (*** p<0.001). (**f**) Distances from selected hotspots to the tumour perimeter, ordered by increasing proximity, across the 12 cases. The dashed line represents no significant difference. The colours depict p-value ranges obtained from Student’s t tests for differences in distance to the tumour perimeter.

To confirm and further explore the emergence of additional cancer hallmarks in the context of EMT, we also defined proliferative, hypoxic and angiogenic hotspots within the same slides (**Fig. 2a**). To check that that SpottedPy accurately measures hotspot distance using the shortest path approach, we simulated a hypoxia hotspot moving away from a mesenchymal hotspot of interest (**Supplementary Fig. 1e**). As the hypoxia hotspot moves farther away from the mesenchymal hotspot the calculated distance between the hotspots increases, as expected.

When visually inspecting the slides, we find angiogenesis and hypoxia frequently accompanying EMT hotspots (**Fig. 2a**). When quantifying hotspot distances using SpottedPy, we confirm that EMT hotspots tend to be closer to angiogenic and hypoxic hotspots compared to EPI hotspots, proliferative hotspots, or the average tumour cell (**Fig. 2b-c**). In contrast, proliferative hotspots were significantly closer to EPI hotspots (p<0.001, **Fig. 2c**). These relationships were consistently observed across breast cancer slides (**Supplementary Fig. 2a**).

To further grasp the positioning of these EMT and EPI areas within the tumour, we used SpottedPy to determine the tumour perimeter (**Fig. 2d**) and calculated distances to it. We conducted a visual benchmark of our tumour perimeter calculation against the Cottrazm method(40), confirming that our algorithm accurately captures similar perimeter trends (**Supplementary Fig. 2b**). We find EMT hotspots closer to the tumour perimeter compared to EPI hotspots, suggesting a state with significant interaction with the surrounding microenvironment (**Fig. 2e**). As expected, angiogenesis hotspots were located closest to the tumour perimeter, followed by hypoxia hotspots (**Fig. 2f**). The spatial localisation of angiogenesis near the tumour perimeter aligns with its function in supplying nutrients and oxygen to rapidly growing tumours(45). The prominence of hypoxic regions succeeding angiogenic zones is consistent with the understanding that rapid tumour growth often outpaces its vascular supply, leading to hypoxic conditions(42). These hypoxic conditions are alleviated in the angiogenic areas, as we find hypoxic coldspots are closest to the tumour perimeter (**Fig. 2f**). Cell proliferation hotspots were observed farthest from the tumour perimeter, and located at spatially distinct locations to EMT hotspots, in line with tumour growth studies outlining a proliferative epithelial core and EMT transformation at the periphery facilitating cancer cell intravasation and migration(46,47).

### EMT hotspots are immunosuppressed and shielded by myCAFs and macrophages

Having confirmed that SpottedPy is able to recapitulate expected spatial hallmarks of EMT within the breast cancer tissue, we next expanded our analysis to dissect the interplay between tumour cells undergoing EMT and other immune and stromal cells in the microenvironment. Alongside EMT hotspots, we calculated hotspots for 41 cell types in the TME, including different lymphocyte, myeloid and fibroblast populations, as defined by Wu et al(12) (**Fig. 3a-c**). When visually inspecting these hotspots, we observed myofibroblastic CAF (myCAF) hotspots and EMT hotspots tended to co-localise (**Fig. 3a-c**). Quantifying hotspot distances using SpottedPy allowed us to confirm that tumour EMT hotspots were indeed closer to myCAF hotspots (**Fig. 3d**). The relationship is particularly highlighted when we look at the cellular niches that are significantly closer to EMT hotspots compared to EPI hotspots, revealing a predominance of various CAF subtypes. This is well in line with existing studies, as myCAFs have been shown to produce TGF-β, which is a well-known EMT trigger(48). They have also been linked to ECM deposition and suppression of antitumor immunity(49),(50),(51),(52).

**Figure 3:**
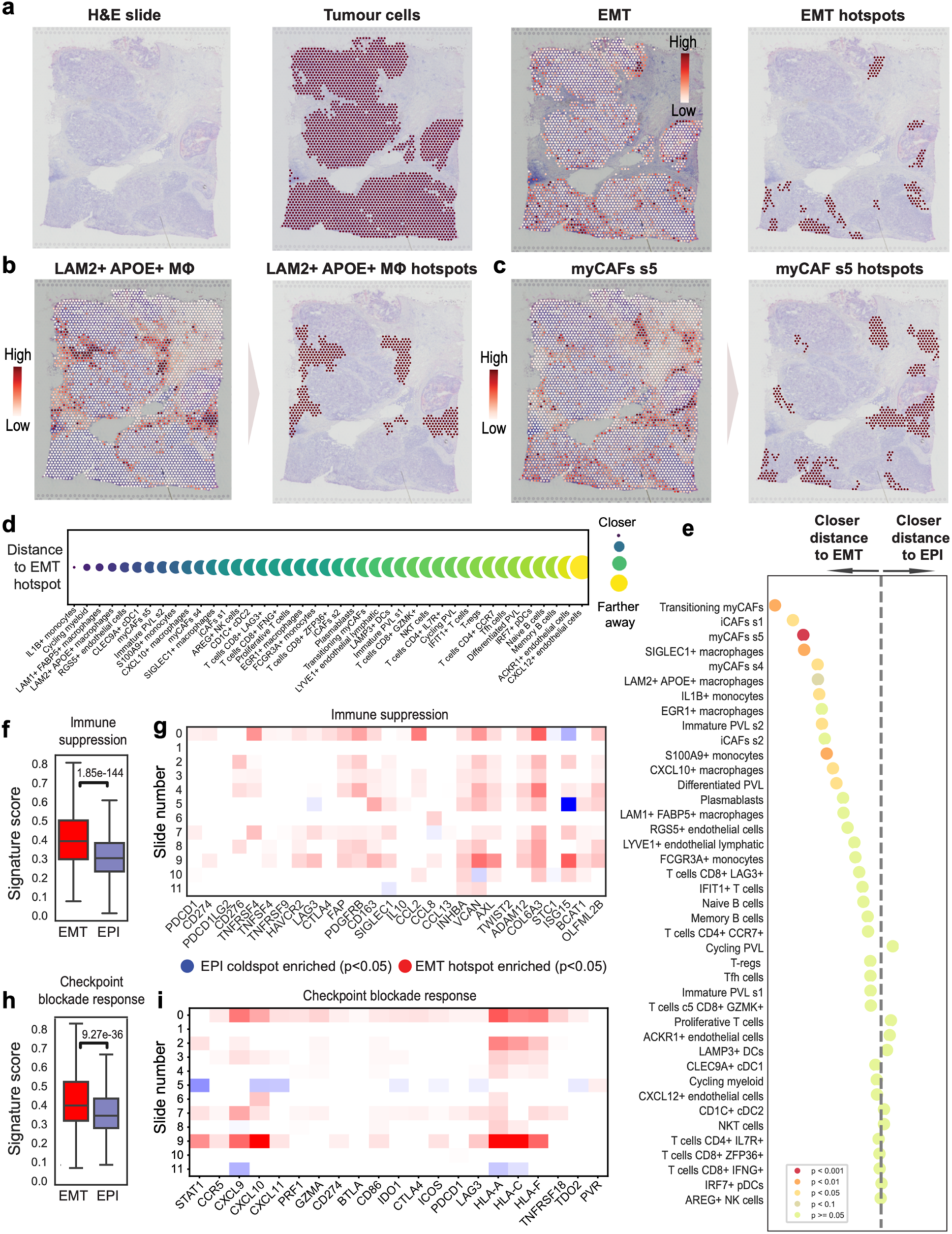
The spatial interplay between EMT progression and the TME. (**a**) Spatial transcriptomics plots highlighting tumour cell spots (left), the EMT gradient through these tumour spots (middle) and EMT hotspots identified by SpottedPy (right) in Slide 5. (**b**) Spatial localisation of macrophage-enriched spots (left) and SpottedPy-defined LAM2+ APOE+ macrophage hotspots (right) in Slide 5. (**c**) Spatial localisation of myCAF s5-enriched spots (left) and SpottedPy-defined myCAF hotspots in Slide 5. (**d**) Distance between EMT hotspots and different TME cell hotspots, ranked by proximity. Smaller, darker bubbles represent shorter distances to EMT hotspots. Results are averaged over 12 slides. (**e**) Distances from various cells in the TME to EMT/EPI hotspots. The dashed line represents no difference in proximity to either EMT hotspots or EPI hotspots. The dots situated to the left of the dashed line indicate cell populations that are significantly closer to EMT hotspots, ordered by decreasing proximity. The colours indicate the p-value ranges obtained from the GEE fit for differences in distance to EMT hot/ EPI hot areas. Results are across 12 slides. (**f**) Barplots showing signature scores of immune suppression scored within EMT hotspots and EPI hotspots(12,55). (**g**) Differences in the average expression of genes in the immune suppression signature between EMT and EPI hotspots for each slide (row). Red depicts genes significantly upregulated in EMT hotspots and blue indicates genes significantly upregulated in EPI hotspots (Student’s t test p<0.05, adjusted for multiple testing using the Bonferroni correction). White indicates a non-significant relationship (12,55). (**h**) Similar to (f) but for checkpoint inhibitor response (61,62) (**i**) Similar to (g) but focusing on the genes in the checkpoint inhibitor response signature.

Additionally, monocytes and particularly tumour-associated macrophages (TAMs) (LAM2+ APOE+ macrophages and SIGLEC1+ macrophages) were prominently closer to EMT hotspots as compared to EPI hotspots. Monocytes, and TAMs derived from them, are known to modulate the environment of tumour cells undergoing EMT, usually by promoting immune suppression in the TME, which facilitates tumour progression and metastasis(53). Natural Killer (NK), NK T cells and CD8+ T cells, the immune cells that can directly kill transformed cells, were amongst the least associated with EMT hotspots, potentially reflecting a mechanism of immune evasion employed by tumour cells that have undergone EMT (**Fig. 3f**). The T-cell sub-population closest to EMT hotspots when compared to EPI were LAG3+ CD8+ T cells, an exhausted population suggestive of immune evasion capacity in these EMT areas (54).

The cohort-level association between EMT hotspots and the myCAF s5 population was maintained in individual tumours, suggesting that this is a universal pattern of EMT transformation in breast cancer and not subtype-specific (**Supplementary Fig. 2a**). In contrast, levels of inter-patient heterogeneity, often even within the same breast cancer subtype, were evident for a variety of cells including macrophages, memory B-cells, naïve B-cells, iCAFs, NK cells, NKT cells, CD4+ T-cells and CD8+ T-cells. However, distances to EMT hotspots were consistent across subgroups of cells (**Supplementary Fig. 2c**), suggesting that, within individual patients, these cells share common response patterns irrespective of the broader heterogeneity observed across the patient cohort.

Due to the close relationship with potential immunosuppressive factors, we next sought to test whether EMT hotspots were indeed likely to be immunosuppressed. We found that EMT hotspots displayed a significantly increased expression of immunosuppression and exhaustion markers (12,55) compared to EPI hotspots (**Fig. 3f-g**), Highly expressed suppressive genes include *FAP*, which has been shown to activate immune suppressive cells such as regulatory T cells (Tregs) and myeloid-derived suppressor cells (MDSCs) (56,57), *INHBA*, which fosters a switch in macrophage polarisation towards a tumour-promoting state (58), *VCAN*, which has been shown to inhibit T-cell proliferation(59) and *COL6A3,* linked to the increased recruitment of macrophages (60). Key immune checkpoints *B7-H3* (*CD276)*, *OX40* (*TNFRSF4*) and *TIM3* (*HAVCR2*) also displayed significantly higher expression (p<0.05) in the tumour slides (**Fig. 3g**). Indeed, EMT hotspots presented increased exhaustion (**Supplementary Fig. 2d-e).** Thus, it appears that the chronic nature of immune activation nearby EMT hotspots leads to the exhaustion of these cells, potentially suggesting opportunities for immune reactivation through checkpoint blockade strategies. To verify this hypothesis, we examined the expression of an interferon-gamma signature that has been associated with response to immunotherapy (61,62) (**Fig. 3h-i**). We found that EMT hotspots had significantly increased expression of genes within the signature, most notably of *HLA-A*,and *HLA-C*, often associated with the activation of immune responses (63). The hotspots also had increased expression of HLA-F, which has immune suppressive functions (64). The expression of interferon-gamma related genes, especially those involved in antigen presentation like HLA molecules, is a favourable prognostic marker in the context of checkpoint blockade therapy (65). These findings suggest that while EMT hotspots are areas of significant immunosuppression and immune cell exhaustion, they also retain elements of immune activity that could be enhanced through targeted therapies such as checkpoint inhibitors.

### EMT hotspots display intra– and inter-patient heterogeneity

We next sought to interrogate spatial relationships at a more granular level, and analysed the association of EMT hotspots with other immune and stromal areas within the same slide and across the different patient samples (**Fig. 4a).** Whilst the cells that had the strongest relationship with EMT hotspots when averaged over the slides, such as SIGLEC+ and LAM2+ APOE+ macrophages and CAFs, displayed the most consistent trends across the slides, it was evident that these relationships were still heterogenous. For example, in slide 4, whilst seven EMT hotspots were closer to LAM2+ APOE+ macrophages than the median EPI hotspots, two were not (**Fig. 4a).** This is further illustrated through visual inspection of the individual EMT hotspots and comparison with the LAM2+ APOE+ macrophage hotspots, enabled through SpottedPy functions (**Fig. 4b).** As expected, stronger associations with myCAFs and macrophage subtypes were pervasive in most tumours and hotspots (**Fig. 4a).** T-cells demonstrated noticeable heterogeneity across patients, but clustered together, reinforcing the idea that these cells share common response patterns. The EMT hotspots that were closer to T-cells were more likely to be enriched in exhaustion markers (**Fig. 4a, right panel),** suggestive of chronic immune activation. We also show that EMT hotspots show a consistent trend of displaying higher suppressive scores than EPI hotspots (**Fig. 4a).** Additionally, nearly all EMT hotspots were closer to the tumour perimeter (**Fig. 4a**), corroborating the overall trends reported in **Fig. 2e-f**. Overall, the inter-patient heterogeneity seemed to supersede the intra-patient heterogeneity.

**Figure 4:**
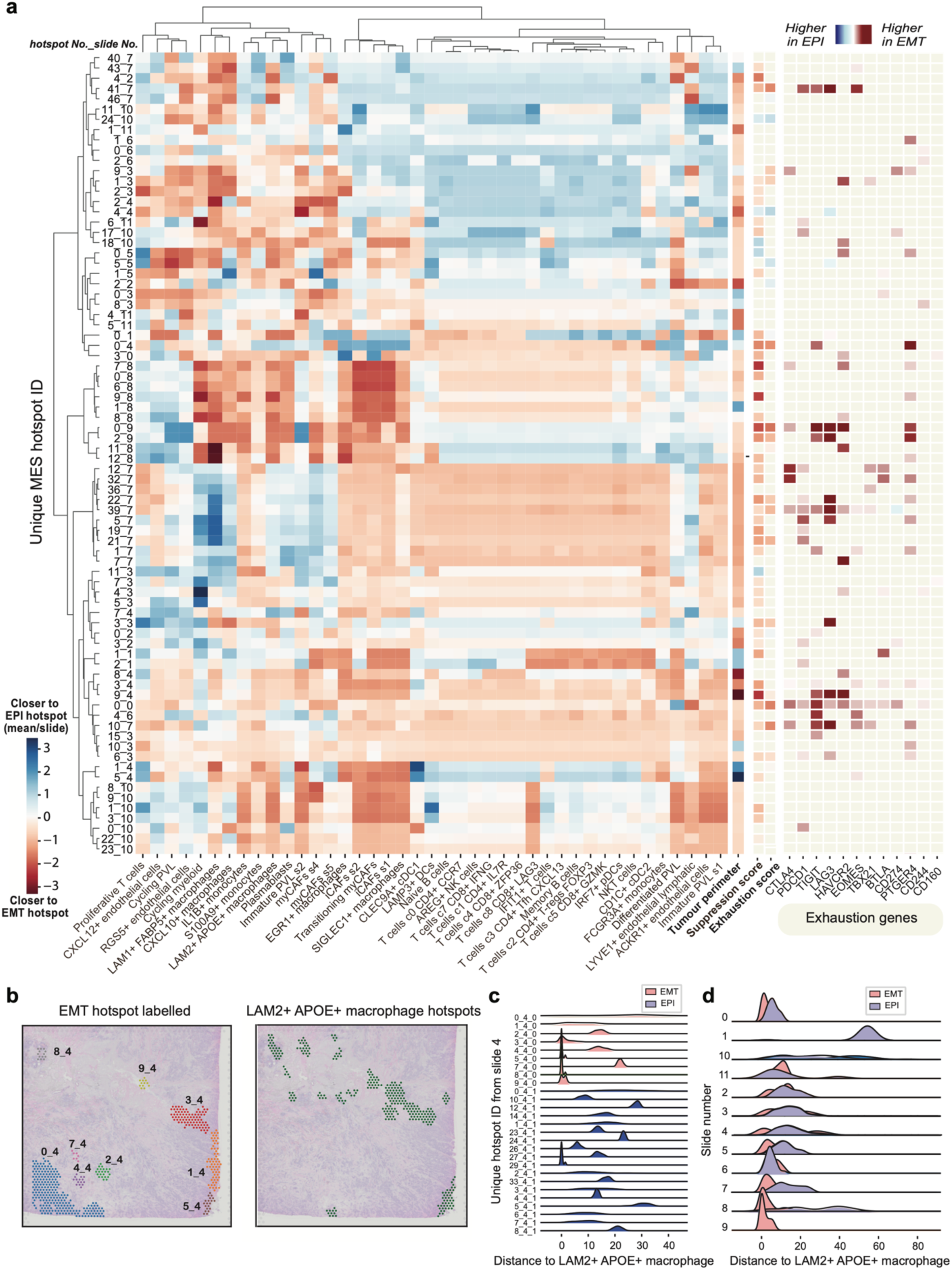
Inter– and intratumour heterogeneity of EMT hotspots. (**a**) Dendrogram highlighting the proximity of EMT and EPI hotspots to TME cell types. The dendrogram is clustered according to the distances from EMT/EPI hotspots to regions enriched in immune/stromal cells. Red indicates that an EMT hotspot is closer to a cell type, while blue suggests that the EPI hotspots in that slide are on average closer. The x-axis displays individual EMT hotspots (label indicates hotspot number and slide number). To the right of the dendrogram, distances to the tumour perimeter, suppression and exhaustion signature scores are illustrated. Red indicates that the hotspot is significantly enriched in these signatures compared to the average EPI hotspot in the slide (p<0.05), blue indicates EPI hotspots are significantly enriched (p<0.05). Further to the right, individual genes associated with the exhaustion signature are shown, with red indicating the gene expression is higher in EMT hotspots (p<0.05) and blue indicating the gene expression is higher in EPI hotspots (p<0.05). (**b**) Slide 4 with individual EMT hotspots labelled (left) and LAM2+ APOE+ macrophage hotspots highlighted (right). (**c**) Distance distributions for each EMT hotspot in slide 4 to LAM2+ APOE+ macrophage hotspots. (**d**) Distance distributions of EMT hotspots to LAM2+ APOE+ macrophages across all 12 slides in the cohort.

SpottedPy offers functionality to inspect the distance distributions for hotspots within a slide (**Fig. 4c),** and across each slide (**Fig. 4d).** Visualising these distributions highlights that whilst LAM2+ APOE+ macrophages are on average closer to EMT hotspots compared to EPI hotspots, there is heterogeneity within each slide and across different slides.

Overall, these results showcase the range of hotspot analyses enabled by the SpottedPy package and the potential to uncover useful biological insights.

### Sensitivity analysis of hotspots

Determining how the hotspot size, governed by the parameter specifying the number of nearest neighbours for the Getis-Ord Gi* metric, affects spatial relationships is crucial for meaningful spatial analysis. We systematically increased hotspot dimensions (**Fig. 5a-b**) to assess the consistency and robustness of identified spatial associations. We find that associations between EMT hotspots, hypoxia and angiogenesis as well as mutual exclusivity with proliferative hotspots are robust and consistent features of the tumour microenvironment, with such relationships remaining remarkably stable across a range of hotspot dimensions (**Fig. 5c**). The cell populations that we previously identified as having the nearest proximity to EMT hotspots at a fixed parameter size (myCAFs, macrophages and monocytes) also maintained this relationship when varying hotspot sizes. Cells that were farther apart presented less stable associations, such as CD8+ LAG3+ T cells where the relationship broke down at a hotspot size of 250, and naive B-cells where the relationship changed multiple times with increasing hotspot size (**Fig. 5c**, **Supplementary Fig. 3a**). These findings suggest that interactions with certain cells in the TME may be more pronounced and relevant at a smaller scale. We found that proliferative hotspots were the most consistently adjacent to EPI hotspots at various hotspot sizes. Adjusting the p-value cut-off used to detect spatial clusters using the Getis-Ord Gi* highlighted similar relationships (**Supplementary Fig. 3b**). To evaluate the stability of the spatial relationships, we introduced gaussian noise and spot reshuffling into the dataset and examined the persistence of these relationships (see Methods). This approach demonstrated that the method is robust to low levels of noise, but also effectively discriminates between biologically meaningful signals from those arising from random spatial distributions (**Fig. 5e-f).** While random noise simulated through spot reshuffling can mimic some aspects of structured data (**Supplementary Fig. 3c**), the hotspots are significantly smaller than those that are biologically relevant (**Supplementary Fig. 3d**). Crucially, the loss of specific associations among particular cell types when noise is introduced (**Fig 5e-f**) contributes to the reduction of false positives even if hotspots are identified(66).

**Figure 5:**
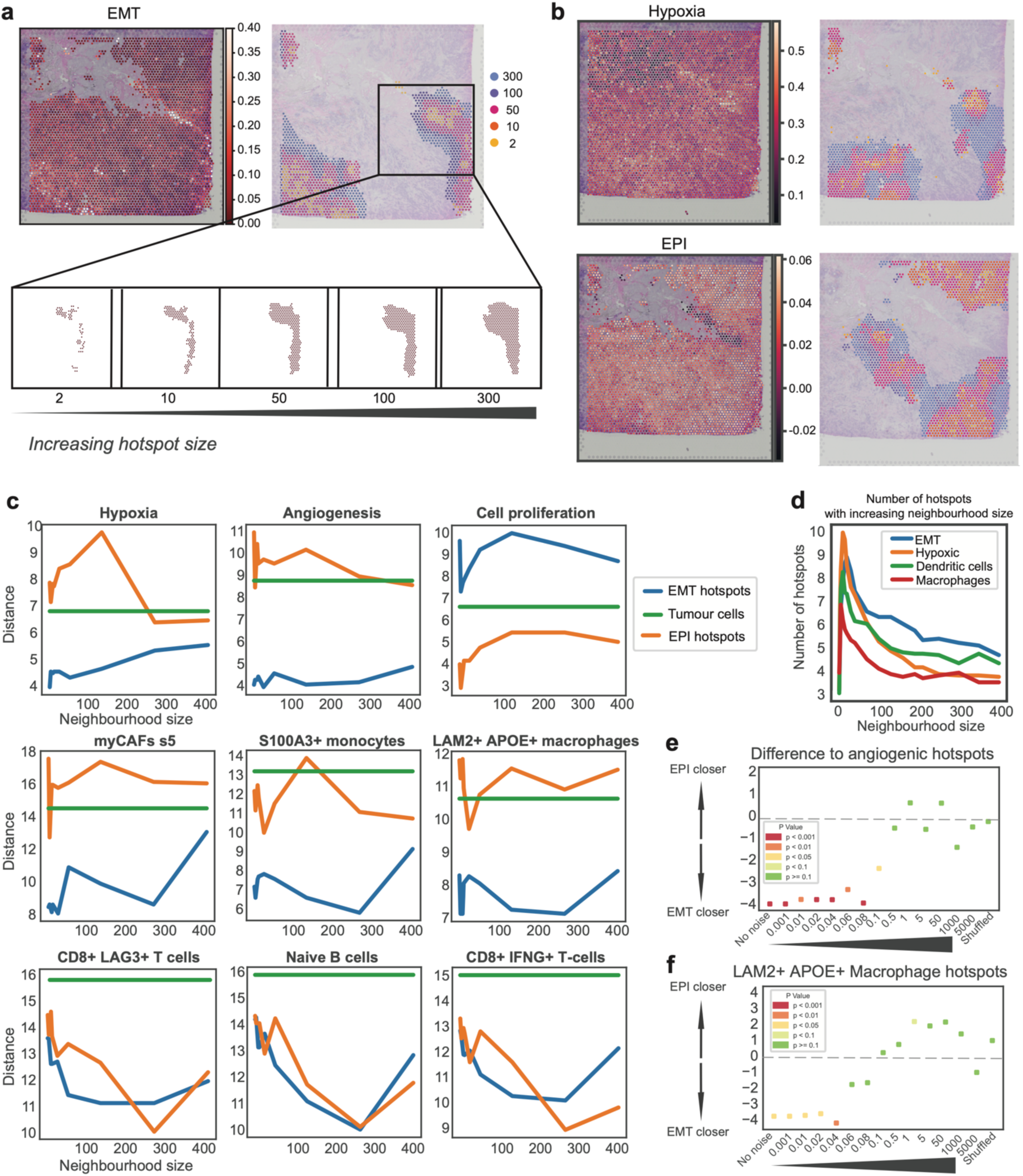
Sensitivity analysis of hotspot relationships. (**a**) EMT hotspot generation using a hotspot neighbourhood parameter of 2, 10, 50, 100 and 300, respectively. Increasingly larger neighbourhoods are highlighted in different colours as indicated in the legend. (**b**) Hypoxia and epithelial hotspot generation using a hotspot neighbourhood parameter of 2, 10, 50, 100 and 300, respectively. (**c**) Sensitivity plots highlighting the distance from EMT hotspots (blue) and EPI hotspots (orange) to regions enriched in various cancer hallmarks and TME components as the hotspot size increases. The distances to the average of all tumour cells are used as a reference (green). Distances are averaged over all 12 slides. (**d**) Number of distinct hotspots identified as the hotspot neighbourhood parameter is increased, averaged over 12 slides. (**e**) Evaluating the impact of increased noise or spot shuffling on the association between angiogenic hotspots and EMT/EPI hotspots. (**f**) Evaluating the impact of increased noise or spot shuffling on the association between LAM2+ APOE+ macrophage hotspots and EMT/EPI hotspots.

### Other distance metrics

We note that there are alternative methods to assess hotspot distances. The “centroid to centroid” methodology offers a practical and straightforward way to approximate the distances between hotspots, but it is crucial to acknowledge its simplicity. As depicted in **Supplementary Fig. 3e**, the size of a hotspot significantly influences its centroid location. Consequently, a larger hotspot, despite being physically closer, may appear farther away when measuring centroid distances, due to its centroid being located farther from the point of interest compared to a smaller, neighbouring hotspot. Thus, the centroid approach may miss local variation that the shortest path method can capture. This could suggest that applying it on our breast cancer slides could potentially miss the more complex relationship observed between EMT hotspots and macrophage or monocyte-enriched areas.

### Spatial EMT relationships in other cancer types

We further investigated whether the relationships observed for EMT hotspots in breast cancer were consistent across other cancer types. We assessed these relationships in publicly available datasets from basal cell carcinomas (BCC)(67), pancreatic ductal adenocarcinomas (PDAC)(68) and colorectal cancers (CRC)(69).

Within the BCC slides, angiogenetic and hypoxic hotspots were closer to EMT hotspots (**Supplementary Fig. 4a-b).** Interestingly, proliferative hotspots were also closer to EMT hotspots, suggesting an alternative relationship compared to breast cancer. POSTN+ fibroblasts were closer to EMT hotspots, whilst there were no significant spatial relationships with T cells or NK cells, paralleling the findings in breast cancer.

We next assessed the relationships within one available PDAC slide (**Supplementary Fig. 4c).** Both angiogenesis and fibroblasts displayed a spatial relationship like that observed with EMT hotspots in breast cancer. In contrast, immune cells were located nearer to EMT coldspots, and there was no significant association between EMT hotspots and hypoxic environments, diverging from the patterns observed in breast cancer.

In CRC, myofibroblasts, angiogenesis, and hypoxia showed comparable spatial relationships to those seen in breast cancer (**Supplementary Fig. 4d-e).** Notably, regulatory T-cells, T-helper cells, and NK cells were significantly closer to EMT hotspots, potentially indicating enhanced immune recognition around EMT hotspots compared to other cancer types.

While limited in breadth and cell type resolution, these analyses suggest that the interplay between tumour cells undergoing EMT and other immune and stromal cells within the TME is likely to be tissue specific, and future work should explore this in more detail.

### Neighbourhood enrichment analysis

The neighbourhood enrichment technique captures more localised, shorter-range relationships with the TME. Additionally, it can assess spatial relationships of phenotypes that would be considered scattered (states that do not occur spatially clustered and therefore might be overlooked by a hotspot-based approach). We experimented with two approaches, ensuring a robust analysis that is less sensitive to the MAUP (**Fig. 1bi-ii**). We first assessed how the spatial relationships change by correlating phenotypes across a central tumour spot and the direct neighbourhood surrounding it (a ring encompassing six Visium spots). We then assessed how the phenotypes are linked within a spot and then expanding what is considered a spatial spot. Varying the method and the number of rings in both cases enables us to assess whether the observed hotspot relationships shift with the unit of analysis and indicates how large of an influence the EMT regions have on surrounding spots.

Our analysis revealed that angiogenesis, myCAFs, macrophages and monocytes exhibited the most significant correlation, in descending order, with cells undergoing EMT (p<0.001) across the 12 slides (**Supplementary Fig. 5a**). This finding reinforces the spatial relationships that we previously identified using the hotspot method. A weaker association was evident for naive B-cells, T-cells, NK cells and NKT cells, in accordance with the hotspot approach. We also found that these spatial relationships were stable across various neighbourhood sizes (**Supplementary Fig 5b**).

The methods show broadly similar trends, suggesting the cellular relationships observed occur both due to colocalization in a spot as well as diffusing influence around the spot.

### EMT state fluctuations shape distinct immune niches within the same tumour

As mentioned previously, EMT is not a binary process – instead, cells are found to occupy multiple hybrid states during the E/M transition. We sought to investigate the spatial distribution of tumour hotspots occupying epithelial (EPI), early intermediate (EM2, EM3), late intermediate quasi-mesenchymal (M1) and fully mesenchymal (M2) states using our multi-scale approach. We captured distinct gene programs representing these states using Non-Negative Matrix Factorisation (NMF) via the CoGAPs workflow(70) (**Supplementary Fig. 6a**). The corresponding hotspots occupied distinct spatial locations within the tissue (**Supplementary Fig. 6b**). When visually inspecting these hotspots, we detected a progressive transformation in the tumour, as highlighted in Slide 4 (**Fig. 6a**). This transition was marked by a spatial shift from EPI into the M1 state, with EM3 serving as an intermediate stage. EM2 displayed volatility in this progression, while M2 was predominantly co-localised with the EPI state. The experimental study by Brown et al(41) had detected that M2 cell clones gained integrin β4 (a key epithelial marker) when cultured, which might have played a significant role in steering these cells towards adopting characteristics more akin to an epithelial phenotype. This would possibly explain the co-localization of these two states within the spatial transcriptomics slide.

**Figure 6:**
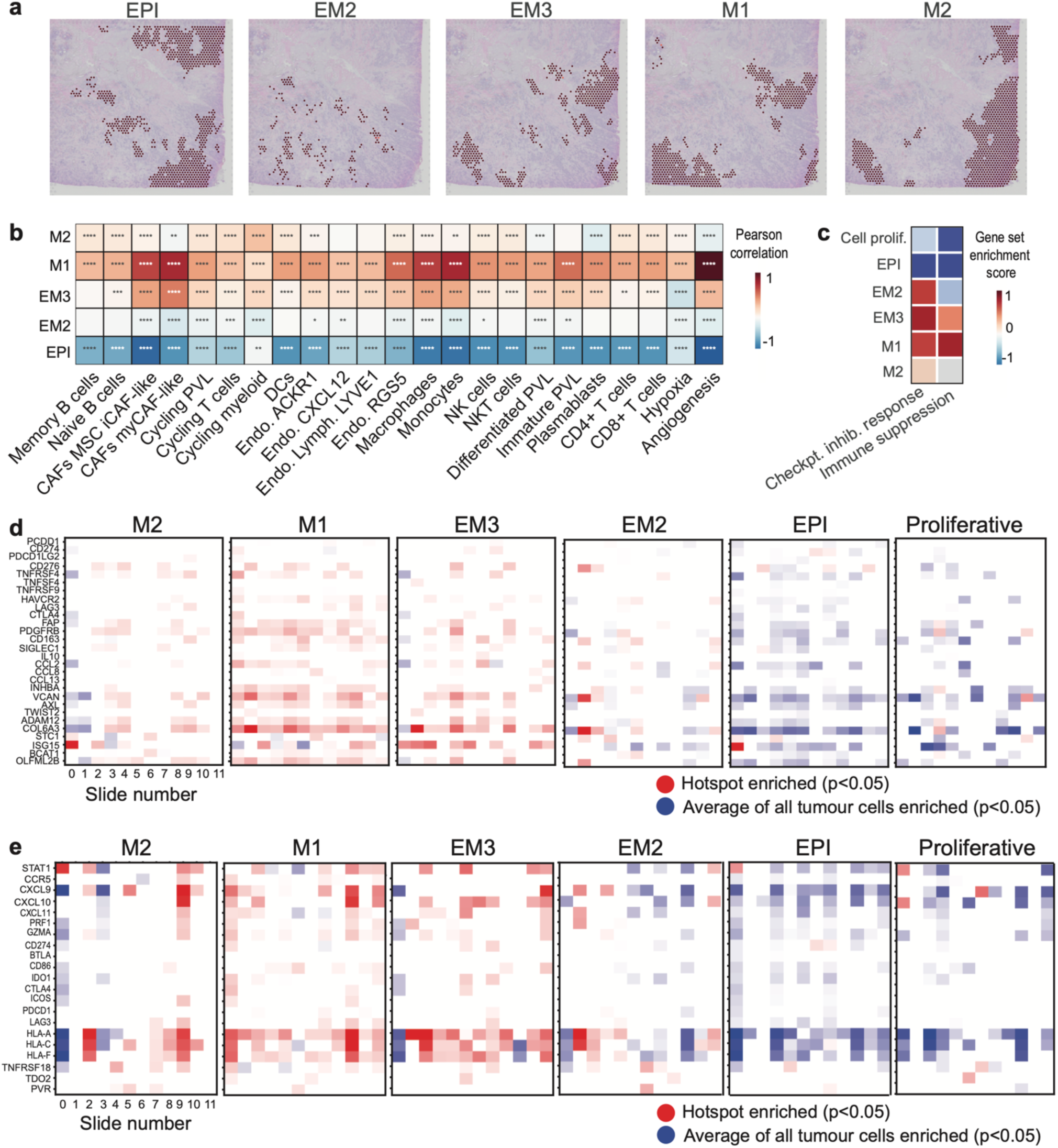
EMT state dynamics uncovered from the spatial exploration of breast cancer tissue. (**a**) Spatial plots depicting epithelial (EPI), intermediate (EM2, EM3), quasi-mesenchymal (M1) and fully mesenchymal (M2) hotspots in Slide 4. (**b**) Neighbourhood enrichment analysis depicting the association between tumour cells occupying distinct EMT states and other cells in the immediate TME, summarised across all 12 slides. Red indicates a significant positive correlation (Pearson, p<0.05), blue a significant negative correlation (p<0.05) and white a non-significant correlation (p>0.05). **** p<0.0001, *** p<0.001, ** p<0.01, * p<0.05. (**c**) Scaled immune suppression(12) and immunotherapy signature(65) scores calculated using Gene Set Enrichment Analysis (GSEA) for each EMT state hotspot and proliferative hotspot, summarised across the 12 samples. (**d**) Enrichment and depletion of expression for genes in the immune suppression signature within EMT state hotspots for each slide (column). Red depicts genes significantly upregulated in EMT state hotspots compared to the average of all tumour cells and blue represents genes significantly downregulated in the EMT state hotspot (Student’s t test p<0.05). White indicates a non-significant relationship. P-values were adjusted for multiple testing using the Bonferroni correction. (**e**) Similar to (d), focusing on genes in the checkpoint inhibitor response signature.

To investigate the relationship between these states, we correlated them with each other and with the generic EMT hallmark signature employed previously for each tumour spot (**Supplementary Fig. 7a**). The lack of significant positive correlation between EPI, EM2, EM3, M1, M2 suggests that these are discrete EMT states. We found a significant correlation between the EMT hallmark signature and the quasi-mesenchymal M1 state, suggesting these are likely capturing a similar state. The EPI state was negatively correlated with the EMT hallmark signature, as expected. EM2, EM3 and M2 are likely distinct states, as evidenced by the lack of correlations across the spots. In terms of spatial distribution, the EMT hallmark hotspots were located closest to the M1 hotspots, and furthest away from the EPI hotspots (**Supplementary Fig. 7b**), in line with the correlation analysis and confirming the hypothesised identities of these states.

When investigating how tumour cells occupying distinct EMT states relate to their microenvironment, we found that the EPI state has negative correlations with immune and stromal cells within the TME, suggestive of a state that is not directly being shaped by the TME (**Fig. 6b**). Interestingly, the M1 state had stronger associations overall with a wide range of cells within the TME, with the strongest relationships established with myCAFs, macrophages and monocytes. We observed similar but weaker correlations with the EM3 state, and considerably weaker correlations with the EM2 state. The progressive loss of association with cells in the EM3 and EM2 states is in line with the idea of these states representing intermediate, more plastic states preceding the apparently more stable M1 state. The M1 state is observed in proximity to natural killer (NK) cells, which is a distinct deviation from the EMT hallmark signature. This observation suggests that while there are resemblances between the M1 and EMT hallmark signatures, the M1 state is likely representative of a unique cellular phenotype with the potential to recruit cells capable of directly eliminating cancer cells.

Furthermore, the quasi-mesenchymal M1 state presented an enrichment of markers linked with immunosuppression and positive response to checkpoint inhibitors (**Fig. 6c**, **Supplementary Fig. 7c**), with a subset of genes, most notably *OX40 (TNFRSF4), TIM3* (*HAVCR2*), *HLA-DRA, CXCL9* and *CXCL10*, driving these relationships (**Fig. 6d-e**). The intermediate states are to a certain extent on the way to adopting this immune suppressive phenotype, with weaker relationships observed with the EM2 state, a slightly stronger enrichment with EM3 and the strongest score with M1 (**Fig. 6c-e**, **Supplementary Fig. 7d**). In contrast, the M2 state has a unique phenotype, and displays both positive and negative relationships with genes within these signatures. We compared these states to the proliferative signature and found that proliferative hotspots mirror the relationship of the EPI state, suggesting that it is a tumour state that is not linked to immunosuppression.

Overall, this analysis sheds light on the changing landscape of tumour-TME interactions during EMT progression in breast cancer, highlighting both intratumour heterogeneity as well as universal interactions that could be exploited for therapy.

## DISCUSSION

In this study we introduce SpottedPy, a Python package that identifies tumour hotspots in spatial transcriptomics slides and explores their interplay with the TME at varying scales. We show that the Getis-Ord G* statistic can be successfully applied to delineate cellular hotspots and provide meaningful biological insights into the spatial organisation of the tumour tissue in its immune and stromal contexture. Whilst various studies have recently applied “hotspot”-type of analysis to spatial transcriptomic data(71),(72),(21), these methods do not offer a way to assess the confidence level in the identification of specific clusters/hotspots, whereas our method assigns a p-value to each hotspot which can be flexibly tuned to according to the user’s stringency requirements. Furthermore, other available methods do not extensively analyse the distances between hotspots. We build on these approaches to analyse the spatial relationships between hotspots in a statistically principled manner, with the additional ability of anchoring hotspot identification to specific regions of interest, such as tumour cells or non-tumour cells, enabling us to interrogate the spatial dynamics within the TME via a more targeted approach. By computing and statistically comparing distances, we offer an interpretable and intuitive measure of relationship between spatial variables. This approach further allows differential spatial analysis between a hotspot of interest and a reference region which other methods do not include. Our method additionally assesses the effect of hotspot size on spatial relationships and compares hotspot spatial trends to the relationships captured using neighbourhood approaches, more frequently applied in spatial transcriptomic analysis, to build layers of spatial evidence.

By adopting our SpottedPy methodology to explore the spatial dynamics of tumour plasticity phenotypes in breast cancer, we have uncovered key differences between tumour regions undergoing EMT and those lacking evidence for this transformation. Our approach illuminates the pronounced spatial correlations of EMT with key cancer hallmarks, notably hypoxia and angiogenesis, in line with findings by He et al(11) in breast cancer spatial transcriptomics detecting these signatures overlapping certain niches. As tumour cells undergo EMT in response to hypoxic stimuli, they are likely to gain a survival advantage in a nutrient-deprived environment and be better equipped to invade and migrate towards regions with better oxygenation, potentially following angiogenic gradients(73),(44),(47).

We find a strong relationship between EMT and CAFs across all slides. CAFs have been linked to tumour cells undergoing EMT in our previous bulk and spatial transcriptomic analyses(33), and have been shown to induce EMT in endometrial cancer cells(74) and hepatocellular carcinoma(75). It is worth noting that CAFs share similar genes with EMT signatures and therefore differentiating between these two cell types can be challenging, notably in bulk tumour settings(76). Here we have leveraged scRNA-seq for deconvolution, alongside detecting copy number aberrations to confirm the presence of tumour cells, which adds a further layer of confidence to the accurate delineation of these cell populations. However, the accuracy of this separation cannot be fully guaranteed and future research investigating this relationship using single-cell resolved spatial transcriptomic data would allow us to confirm this relationship more confidently. We further note that while we have tried to ensure EMT is captured only within the tumour cells themselves by investigating signatures only within the tumour spots identified by STARCH and by using EMT reference datasets from pure tumour populations, further validation of tumour regions, e.g. by staining with specific markers, would be beneficial for users who wish to employ our method in their spatial transcriptomics experiments.

In addition to expected CAF associations, we observe a strong relationship with macrophages and monocytes across multiple spatial scales. We particularly observed relationships with SIGLEC+ macrophages, LAM2+ APOE macrophages and EGR1+ macrophages, which are analogous to M2-like, tumour promoting, macrophages(12). Macrophages secrete TGF-β, TNF-α, IL-6 and IL-8, which are well-characterised EMT stimuli(77),(78). The relationship has been observed in bulk transcriptomics(79), in spatial analysis of mouse models of skin carcinoma where depletion of macrophages inhibited EMT progression(30) and in specific niches within breast cancer spatial transcriptomics slides(11). Whilst the relationships with CAFs and macrophages were displayed across the majority of tumour slides, within each slide there were EMT hotspots where this relationship was less clear. This indicates that other factors which we have not accounted in our analyses could play a significant role in driving EMT within local niches.

We uncovered heterogeneous relationships across breast tissue slides with other key immune cells such as NK cells, NKT cells and T cells, highlighting the multifaceted interplay between these components. T-cells have been shown to induce EMT in breast cancer(80),(81), and this relationship has further been highlighted in bulk transcriptomics(82) and smaller scale spatial transcriptomic analyses(33),(83) however there is also evidence showing the exclusion of these cells, linked to the relationship between EMT, macrophages and CAFs promoting an immune suppressed environment(77),(52),(84). Indeed, our findings uncover notable associations between EMT hotspots and immune suppression, alongside signatures indicative of a response to checkpoint therapy, building upon evidence that EMT may offer crucial insights for existing strategies in immunotherapy(85),(11).

Our analysis reveals that EMT occurs in discrete spatial locations distinct from proliferative signatures. This finding is in line with a recent analysis of breast cancer by Jia et al^51^ utilising a more focused spatial transcriptomic dataset, previous research by Tsai et al(86) demonstrating that a departure from a mesenchymal-like state is a prerequisite for tumour cell proliferation in mouse models, and a recent study by Chen et al(87) investigating EMT states in scRNA-seq data. Such spatial characterisations at various scales were largely unexplored.

Delving deeper into EMT, we observed that hybrid EMT states exhibit more heterogeneous and weaker associations with tumour-promoting populations in the TME in comparison to the quasi-mesenchymal M1 state. This disparity might be indicative of the inherent plasticity of hybrid EMT states(88),(89), complicating our ability to delineate clear relationships, but might also suggest a directed trajectory towards an M1 state. The M2 state demonstrates more similar distribution and TME associations to the EPI state, which may be attributed to the activation of integrin β4 (a key epithelial marker) when cultured, a limitation mentioned in the original study which potentially transformed the state towards a more epithelial phenotype(41).

These results point to a highly dynamic and plastic nature of tumour cells in navigating the complexities of their microenvironment. The interactions likely extend beyond a linear framework. Hypoxia, a known catalyst for both angiogenesis and EMT(90), can set off a cascade of events that not only amplify these processes but also create a conducive environment for the recruitment of immunosuppressive cells such as macrophages(91),(92). These cells can in turn bolster angiogenesis, thereby fuelling a self-perpetuating cycle that further complicates the tumour landscape. These insights provide a further understanding of the cellular interactions and environmental factors that underpin tumour progression and metastasis, and could in the future pave the way for the development of targeted interventions aimed at disrupting these complex networks for therapeutic benefit.

The consistency of the relationships we observed across different hotspot sizes and neighbourhood scales further strengthens our confidence in the findings. The neighbourhood ring approach predominantly captured TME cells that have infiltrated the tumour, offering insights into the immediate cellular interplay at the tumour periphery. In contrast, the hotspot methodology provided a broader view, encompassing interactions at more distal locations. By pinpointing statistically significant cellular hotspots, we bolster the reliability of our observations, especially considering the inherent inaccuracies that can arise from deconvolution algorithms applied to non-single cell transcriptomic datasets such as those from the Visium platform.

To the best of our knowledge there are no direct comparisons available for our SpottedPy methodology due to the unique nature of focusing on discrete spatial clusters of user-defined continuous signatures at expanding scales, and performing differential spatial relationships compared to a reference for downstream analysis.

Our insights into the spatial organisation of tumours during EMT progression are limited by the significant amount of uncertainty surrounding EMP programmes (31),(32) and their incomplete characterisation in different types of breast cancer and other neoplasms. In the future, integrating further hybrid states characterised in other breast cancer studies(87) will help expand our understanding of this complex process alongside its multiple locally stable peaks and valleys. The heterogeneous relationships observed with CD8+ and CD4+ T-cells and NK cells require further experimental validation and exploration. As spatial transcriptomics datasets become more widely available, expanding this analysis beyond the current 12 slides could help clarify the perceived spatial heterogeneity and better distinguish universal relationships from local, patient-specific effects.

When investigating to what extent these spatial EMT relationships are maintained or differ across cancer types other than breast cancer, we were limited by the availability and size of such datasets, as well as the differences in cell composition between tissues. Ultimately, any uncovered differences are likely attributed to the unique TME and genetic basis of each cancer type, and in the future a more in-depth analysis in larger datasets once these become more widely available will shed light on the heterogeneity of these relationships. Additionally, extending these analyses to single cell-resolved spatial datasets and incorporating ligand-receptor signalling information into the evaluation of spatial effects on cell populations will increase the confidence in the identified relationships.

Overall, our findings confirm expected spatial effects of EMT progression in tumours, demonstrating that SpottedPy can capture complex associations between tumour cells and their microenvironment. Such insights can help unveil local effects of the TME and linked tumour cell vulnerabilities that could ultimately be exploited for therapeutic benefit. While the analyses presented here primarily illustrate insights into breast cancer tissue organisation, we note that SpottedPy can be applied to discern spatial relationships in other cancer types (as briefly demonstrated) as well as other diseases and even within healthy tissue. SpottedPy has been developed on spatial transcriptomics data from the 10x Visium platform; however, we note it can be easily extended to other spatially-resolved platforms and future releases will provide further functionality to enable this.

## Conclusion

In conclusion, SpottedPy provides a detailed and multifaceted analysis of the spatial dynamics within individual spatially-profiled tumours. By rigorously investigating the proximities of various cellular components, we have underscored the significant influence exerted by cells undergoing EMT in sculpting the TME and highlight SpottedPy as a package that can be applied to answer other spatial biology questions.

## METHODS

### The SpottedPy package

SpottedPy package is compatible with Python 3.9 and depends on scanpy, libpysal, esda packages.

- GitHub: https://github.com/secrierlab/SpottedPy
- Tutorials: spottedpy_multiple_slides.ipynb (this tutorial walks through using SpottedPy with multiple spatial slides, highly recommended for downstream statistical analysis). spottedpy_tutorial_sample_dataset.ipynb tutorial walks through using SpottedPy with a single slide.
- Sample data: https://zenodo.org/records/10392317 The key functions are outlined in the relevant sections below.

### Spatial transcriptomic datasets

Breast cancer Visium slides were obtained from Barkley et al.(28) (slides 0-2), from 10x Genomics (slides 3-5)(34) and Wu et al.(12) (slides 6-12). Slide annotations, if available, are displayed in **Supplementary Fig. 1b**. We combined the three datasets of breasts cancer 10X Genomics Visium spatial transcriptomic datasets into a common *anndata* Python format for analysis. Pre-processing and normalisation were conducted using the ScanPy (Single-Cell Analysis in Python) package(93). We analysed a total of 32,845 spatially profiled spots, and retained spots if they exhibited at least 100 genes with at least 1 count in a cell, had more than 250 counts per spot and less than 20% of total counts for a cell which are mitochondrial. Pre-processed BCC slides were obtained from Gania et al.(67), PDAC slides obtained from Ma et al.(68) and CRC slides obtained from Valdeolivas et al.(69) We used the deconvolution results provided in each of the source studies.

### Spatial data deconvolution

Due to the imperfect near-single cell resolution of current spatial transcriptomic methods, we require a method to deconvolve each spot in order to infer the cellular populations enriched in each spot. Cellular deconvolution was carried out using Cell2location(35). Cell2location decomposes the spatial count matrix into a predefined set of reference cell signatures by modelling the spatial matrix as a negative binomial distribution, given an unobserved gene expression level rate and gene– and batch-specific over-dispersion. A scRNA-seq breast cancer dataset containing 100,064 cells from 26 patients and 21 cell types from Wu et al(12) was chosen to perform the deconvolution. Cell types in the chosen breast dataset consisted of cancer epithelial cells (basal, cycling, Her2, LumA, LumB), naïve and memory B cells, myCAF-like and iCAF-like cancer-associated fibroblasts, perivascular-like cells (PVL), including immature, cycling and differentiated, cycling T-cells, cycling myeloid cells, dendritic cells (DCs), endothelial cells expressing *ACKR1*, *CXCL12* or *RGS5*, endothelial lymphatic *LYVE1*-expressing cells, luminal progenitors and mature luminal cells, macrophages, monocytes, myoepithelial cells, natural killer (NK) cells, natural killer T (NKT) cells, plasmablasts, CD4+ T cells, and CD8+ T cells. We scored the scRNA-seq cancer epithelial cells with EPI and EMT signatures(36,37), and used Gaussian mixture modelling to assign the cells to EPI and EMT clusters.

The scRNA regression model was trained with 500 epochs, and the spatial transcriptomic model trained with 20,000 epochs using a GPU. To delineate the tumour cells within our spatial transcriptomics dataset, we used the STARCH Python package designed to infer copy number alterations (CNAs)(38). STARCH identifies tumour clones (setting *K*=2 clones) and non-tumour spots. It confirms identification of normal spots by clustering the first principal component into two clusters using K-means. Changing the value of *K* alters the number of identified tumour clones, but the number of cells labelled as tumour cells remains the same. This approach is based on the principle that the direction of maximum variance in the expression data typically reflects the division between non-cancerous and cancerous spots. <u>Only tumour cell spots were considered for EMT analysis. The EPI and EMT spots identified using Cell2location were used to define the EPI and EMT hotspots in the breast cancer downstream analysis</u>.

### EMT state and hallmark signature scoring

To identify more granular distinct EMT states, we employed data from Brown et al(41), consisting of seven RNA-seq sequenced cell clones, derived from SUM149PT inflammatory breast cancer cell line with 3 repeats spanning the EMT spectrum from epithelial-like (EPI), quasi-mesenchymal (M1), fully mesenchymal (M2) and three distinct intermediates (EM1, EM2, EM3). We used these data to derive a weighted gene signature to represent the EMT states. We captured EMT gene patterns from this data using non-negative matrix factorisation (NMF) by applying the CoGAPs workflow(70). We used ProjectR’s implementation of *lmfit* R function to map the captured EMT patterns onto the spatial transcriptomic spots(94). This transfer learning approach assumes that if datasets share common latent spaces, a feature mapping exists between them and can measure the extent of relationships between the datasets. The final states were captured with 20 patterns and 10,000 training iterations. The number of patterns were chosen based on capturing the discrete states with the highest accuracy. The EM1 state was not distinguishable from the EPI state, so we merged the two states. Thus, overall we obtained scores for one epithelial, two intermediate, a quasi-mesenchymal and a fully mesenchymal state for each spot.

Hypoxia and angiogenesis were defined based on signatures deposited at MSigDB(95). The proliferative signature was compiled from Nielsen at al(96). The immunosuppresion signature was compiled from Wu et al(12) and Cui et al(55). The checkpoint blockade response signature was compiled from Johnson et al(61) and Liu et al(62). The exhaustion signature comprised classical exhaustion markers: *CTLA4, PDCD1,TIGIT, LAG3, HAVCR2, EOEMT, TBX21, BTLA, CD274, PTGER4, CD244* and *CD160*(97). All these signatures were scored using *scanpy.tl.score_genes* function. EMT hotspots and coldspots were identified in the BCC, CRC and PDAC slides using the EMT hallmark signature(95).

### Graph construction

The SquidPy(23) (Spatial Single-Cell Analysis in Python) package was used for graph construction using *sq.gr.spatial_neighbors* and slide visualisation of the Visium spatial slides. NetworkX(98) was used for further analysis of the networks derived from the spatial transcriptomic spots. The deconvolved spot results were used to assign node labels. Edges were assigned based on the spot neighbours.

### Neighbourhood enrichment analysis

We calculated neighbours for each spot by summing the deconvolution results in a ring surrounding the spot of interest, and normalising by the number of spots assigned as a neighbour, using the adjacency matrix of the graph to calculate the interacting cells.

Two methods were developed to assess neighbourhood enrichment. *Inner outer* correlation (with the function *sp.calculate_inner_outer_correlations*) was calculated by correlating signatures across a central spot of interest and the direct neighbourhood of spots surrounding it (a ring encompassing six Visium spots), after filtering for tumour spots only. To perform the sensitivity analysis, we increased the number of rings surrounding a spatial transcriptomic spot (setting *rings_range* parameter in *sp.calculate_inner_outer_correlations* function) to consider as spot neighbours and compared the change in correlation coefficient. The first ring consists of 6 spots, and the second ring includes 18 spots (combined from the 1st and 2nd rings). Subsequent rings follow this pattern. The number of rings selected for sensitivity analysis reflects a balance between spatial coverage and resolution. Using a smaller number of rings (e.g., 1, 2, 3) allows the analysis to focus on the immediate microenvironment around the central spot, providing high resolution. As more rings are added, the spatial coverage increases, capturing broader interactions but potentially diluting local-specific signals. Correlations were calculated using Pearson’s correlation coefficient.

An *all-in-one* correlation (*sp.calculate_neighbourhood_correlation* function) was calculated by correlating phenotypes with cells within a spot, and then incrementally increasing the number of rings to correlate across progressively larger spatial units. The functions *sp.correlation_heatmap_neighbourhood* and *sp.plot_overall_change* plot the neighbourhood results.

### Hotspot analysis

Hotspots were calculated using The Getis-Ord G* statistic as implemented using the PySAL package(99), using 10 as neighbourhood size parameter by default (number of spot neighbours surrounding the central Visium spot) and a p-value of 0.05, unless otherwise stated.

The Getis-Ord G* equation is defined as follows:

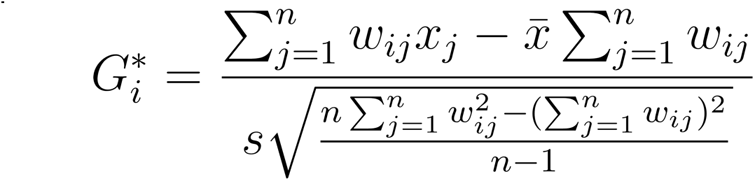

Where *w*_*ij*_ is the spatial weight between location and *i*, 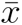 is the mean of the variable of interest across all locations, *s* is the standard deviation of the variable of interest across all locations and *n* is the total number of locations.

A high positive value at location *i* suggests a hotspot for the attribute, while a negative value indicates a coldspot. The significance of *G*\* is determined by comparing the observed 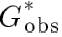 to a distribution of *G*\* values generated under the assumption of spatial randomness. This distribution is obtained by permuting the attribute values across locations and recalculating *G*\* for each permutation. The p-value for a hotspot (when *G*\* is positive) or a coldspot (when *G*\* is negative) is then derived from this distribution. This approach provides a non-parametric method to evaluate the significance of spatial clusters, offering a robust measure against potential spatial randomness in the data. The significance of *G*\* is determined by comparing the observed 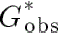 to a distribution of *G*\* values generated under the assumption of spatial randomness. This distribution is obtained by permuting the attribute values across locations and recalculating *G*\* for each permutation. The p-value for a hotspot (when *G*\* is positive) or a coldspot (when *G*\* is negative) is then derived from this distribution.

Hotspots can be identified by calling *sp.create_hotspots* function, and specifying in the *filter_columns* parameter what region within the spatial slide to calculate the hotspot from e.g. tumour cells. The *neighourhood*_*parameter* can be altered here (default = 10). We encourage the user to choose the parameter most relevant for their biological question, e.g. whether they are interested in local interactions of the signature, or broader tissue modules. SpottedPy allows the user to perform the sensitivity analysis to observe how the parameters affect downstream analysis. For the 10x Visium platform, we would recommend starting with parameter k=10 as this captures all the spots surrounding the central spot. The variable with the most stable relationships across a range of parameters (and therefore scales) is likely one of most interest for further investigation. However, specific short-range relationships defined locally rather than across scales could also be of interest in certain circumstances depending on the user’s biological questions. <u>Coldspots are automatically created</u> <u>when *sp.create_hotspots* is called, and hotspots are labelled in the *anndata* object by appending</u> *<u>“_hot”,</u>* <u>and coldspots by appending *“_cold”* to the original column name.</u> <u>When an appropriate</u> <u>contrasting signature is available for comparison e.g. EPI compared to EMT we do not need to use</u> <u>the coldspots for comparison.</u> The *relative_to_batch* parameter ensures hotspots are calculated across each slide, otherwise they are calculated across multiple slides. Importantly, if multiple slides are used (highly recommended for statistical power), these should be labelled using *.obs[‘batch’]* within the *anndata* object. Additionally, the library ID in the *.uns* data slot should be labelled with the*.obs[‘batch’]* value. Hotspots can be plotted using *sp.plot_hotspots*.

Hotspots and coldspots for EMT states and cell proliferation were calculated after filtering for tumour cells as labelled by STARCH, as we aimed to specifically capture these processes within the tumour cells themselves. <u>EMT hotspots are the regions with a high proportion of mesenchymal tumour cells</u> <u>within the tumour-labelled spots. Therefore, they only include a subset of tumour spots. Similarly, cell</u> <u>proliferation hotspots are regions with high fractions of proliferating tumour cells.</u> All other hotspots (deconvolved cell proportion data and angiogenic and hypoxia signatures) were calculated using all the spots within the spatial transcriptomic slide.

### Distance metrics

After calculating the hotspots and coldspots, we then assessed the distances from hotspots of interest (EPI and EMT) to other cells types and signature hotspots and coldspots. We used the ***shortest path*** approach to calculate distances between hotspots as follows:

● Let *H* represent the set of coordinates of spots in the hypoxia hotspot.
● Let *M* represent the set of coordinates of spots in the mesenchymal tumour hotspot.
● Let *E* represent the set of coordinates of spots in the epithelial tumour hotspot.

For a spot *m* in *M* and a spot *e* in *E*, the shortest path to any point *h* in *H* was determined:

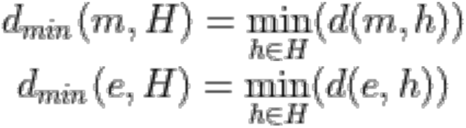

Where *d*(*m*, ℎ) represents the Euclidean distances from a spot *m* in *M*. After obtaining the minimum distances for each spot in *M* and *E* we calculated the median (with the additional functionality to choose min or mean) to provide a summary statistic that reflects the geneal proximity of each hotspot (*M* and *E*) to *H*. The function *sp.calculateDistances* calculates this.

To then infer the impact of cellular hotspots on distance to EMT compared to EPI hotspots, we employed Generalised Estimating Equations (GEE). This model enables us to estimate population-average effects involving repeated measurements across multiple spatial transcriptomic slides. The model estimates the coefficient (β*_mes_*) for the transition from reference hotspots (*E*) to primary hotspot variables (*M*). A positive β*_mes_* would indicate that mesenchymal hotspots are, on average, located further from hypoxic areas compared to epithelial hotspots, while a negative value suggests a closer proximity*. sp.plot_custom_scatter*, setting *compare_distance_metric* to *min, mean* or *median* to compare the summary statistics for each hotspot across each slide. Setting it to *None* calculates the statistical significance of all distances from each hotspot.

The ***centroid approach*** is calculated as follows. The centroid *C*_*H*_ of a set of spots *H* with coordinates *x*_*h*_, *y*_*h*_ is the arithmetic mean of the coordinates. This point represents the centre of the mass of the points in the set *H*.

For set *H*:

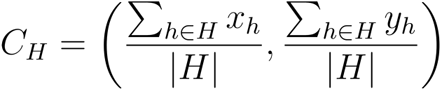

Similar calculations are employed for *M* and *E*. We then calculated the Euclidian distance between the centroids.

### Tumour perimeter calculation

Any spot was considered part of the tumour perimeter if it had more than one neighbouring spots (nodes in the graph) that were not classified as tumour spots. A spot *s* ϵ *S* is considered part of the tumour perimeter, P, if:

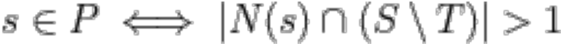

Where *S* denotes the set of all spots, *T* denotes the set of tumour spots N(*s*) represents the neighbouring spots of spot *s*. Additionally, we applied a filtering step to remove isolated perimeter spots. This involved eliminating any identified perimeter spots that had no neighbouring perimeter spots, thereby excluding isolated perimeter spots caused by a single non-tumour labelled spot within the tumour. This approach helped us to delineate the boundary of the tumour accurately by focusing on the transitional area where tumour and non-tumour spots meet (called using *sp.calculate_tumour_perimeter*).

To quantify the number of tumour hotspots, we calculated the number of connected components within the graph that were labelled as hotspots. This calculation was crucial for understanding the distribution and clustering of tumour cells.

### Sensitivity analysis

The sensitivity analysis to evaluate the impact of varying hotspot sizes on the spatial relationships was achieved by incrementally adjusting the neighbourhood parameter for the Getis-Ord statistic, which directly influenced the size of identified hotspots (*sp.sensitivity_calcs)*. As we expanded the neighbourhood parameter, we compared the distances between the newly defined hotspots and other existing hotspots of interest.

To assess the robustness of the spatial relationships between cell types and gene signatures, we systematically introduced Gaussian noise into our cell type proportion data and gene signature matrix. Gaussian noise, characterised by a mean of zero and varying standard deviations, was added to mimic experimental and technical variability. This approach allows us to evaluate the stability of detected EMT hotspots under different noise conditions. We defined a range of sigma values to represent varying levels of noise intensity. To further test the robustness of the spatial relationships, we randomly shuffled the cell proportion data and gene signature values and assessed how this affected downstream analysis.

### Statistical analysis

Groups were compared using a two-sided Student’s t test. Multiple testing correction was performed where appropriate using the Bonferroni method. Graphs were generated using the seaborn and Matplotlib Python packages.

## DECLARATIONS

### Ethics approval and consent to participate

All datasets employed in this study are publicly available and comply with ethical regulations, with approval and informed consent for collection and sharing already obtained by the relevant organizations.

### Consent for publication

Not applicable.

### Availability of data and materials

Breast cancer Visium slides were obtained from Barkley et al. (28), Wu et al. (12) and the 10x Genomics website (34).

SpottedPy is implemented as a Python package available at https://github.com/secrierlab/SpottedPy, released under a GNU General Public License v3.0 with an accompanying tutorial. Processed spatial transcriptomics data have been deposited at Zenodo: https://zenodo.org/records/13284570.

## Author contributions

MS designed the study and supervised the analyses. EW developed the SpottedPy package and performed all the analyses.

