## Supplementary Material for "SpottedPy quantifies relationships between spatial transcriptomic hotspots and uncovers new environmental cues of epithelial-mesesenchymal plasticity in breast cancer"

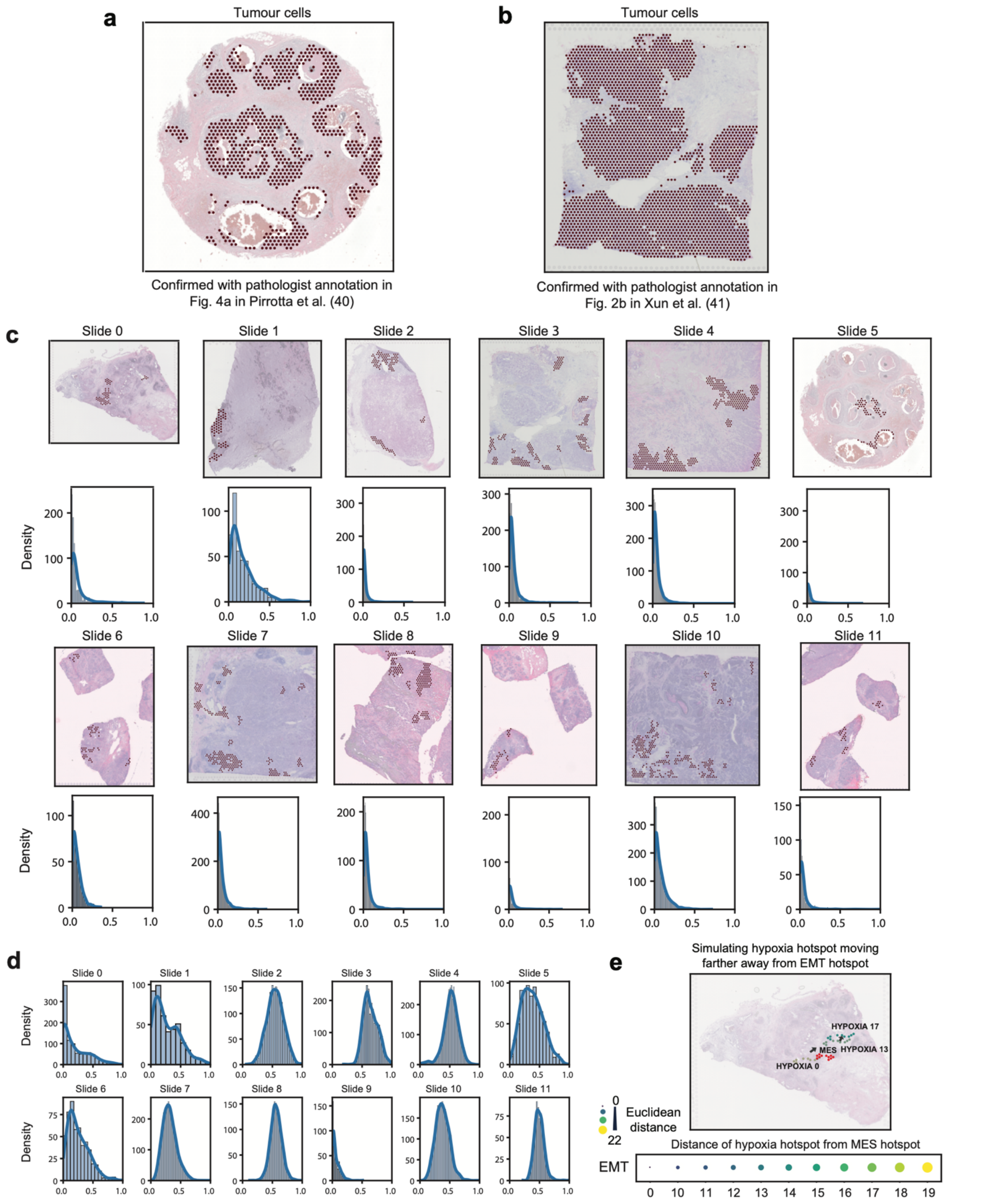


**Supplementary Figure 1: Validation of methods and EMT hotspot distribution.** **(a)** Tumour cells as estimated by STARCH for slide 5, and the reference for the pathologist annotation confirming tumour cell estimation is provided. **(b)** Similar to (b) but for Slide 3. **(c)** Spatial plot depicting EMT hotspots (top) and distributions of EMT fractions (bottom) across slides as returned by Cell2location. **(d)** Distributions of the EPI proportions across slides as returned by Cell2location. **(e)** Spatial plot illustrating simulating a hypoxia hotspot moving further away from an EMT hotspot (top) and the resulting increase in distance returned from SpottedPy calculations (bottom).

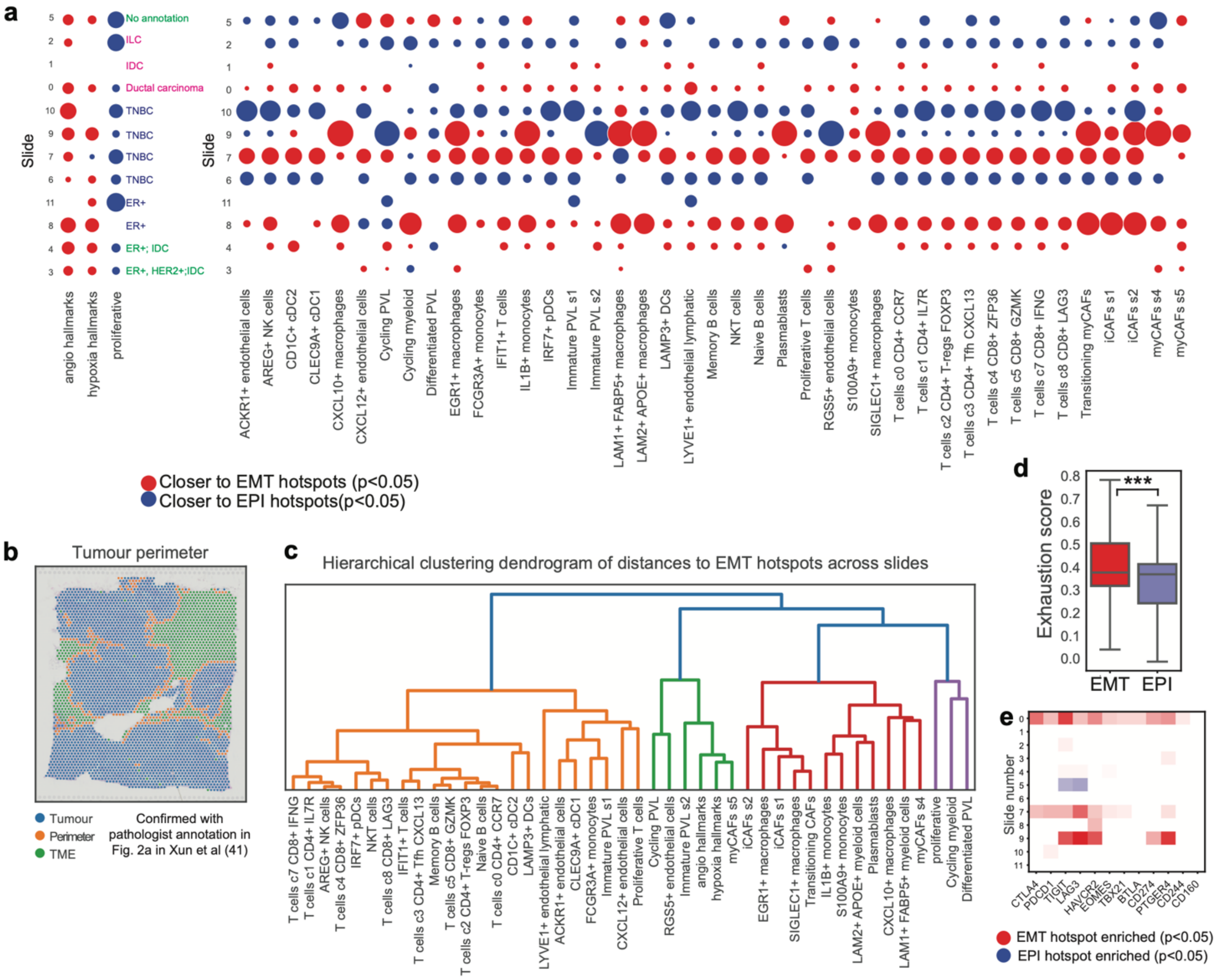


**Supplementary Figure 2: Distribution of cancer hallmarks and cells in the TME in relation to EMT/EPI hotspots by tumour slide. (a)** Bubble plot depicting distances between cancer hallmark signatures and TME classes and EMT/EPI hotspots for each slide (row). Blue depicts hallmarks that are significantly closer to EPI hotspots and red represents hallmarks that are significantly closer to EMT hotspots (Student's t test $p<0.05$), adjusted for multiple testing using the Bonferroni correction. White indicates a non-significant relationship. Tissue annotations, if available, are included on the right-hand side for each sample, coloured by batch. IDC= Invasive Ductal Carcinoma. ILC= Invasive Lobular Carcinoma. **(b)** Tumour perimeter calculated by SpottedPy for slide 3 and the reference for the perimeter confirmed by a pathologist on the same slide. **(c)** A dendrogram illustrating the hierarchical clustering of cells based on their proximity to EMT hotspots, as derived from the data shown in panel (a)**. (d)** Exhaustion signature compared between EMT hotspots and EPI hotspots. **(e)** Expression of individual immune exhaustion genes compared between EMT hotspots and EPI hotspots for each slide (row). Red depicts genes significantly upregulated in EMT hotspots and blue indicates genes significantly upregulated in EPI hotspots (Student's t test $p<0.05$, adjusted for multiple testing using the Bonferroni correction). White indicates a non-significant relationship.

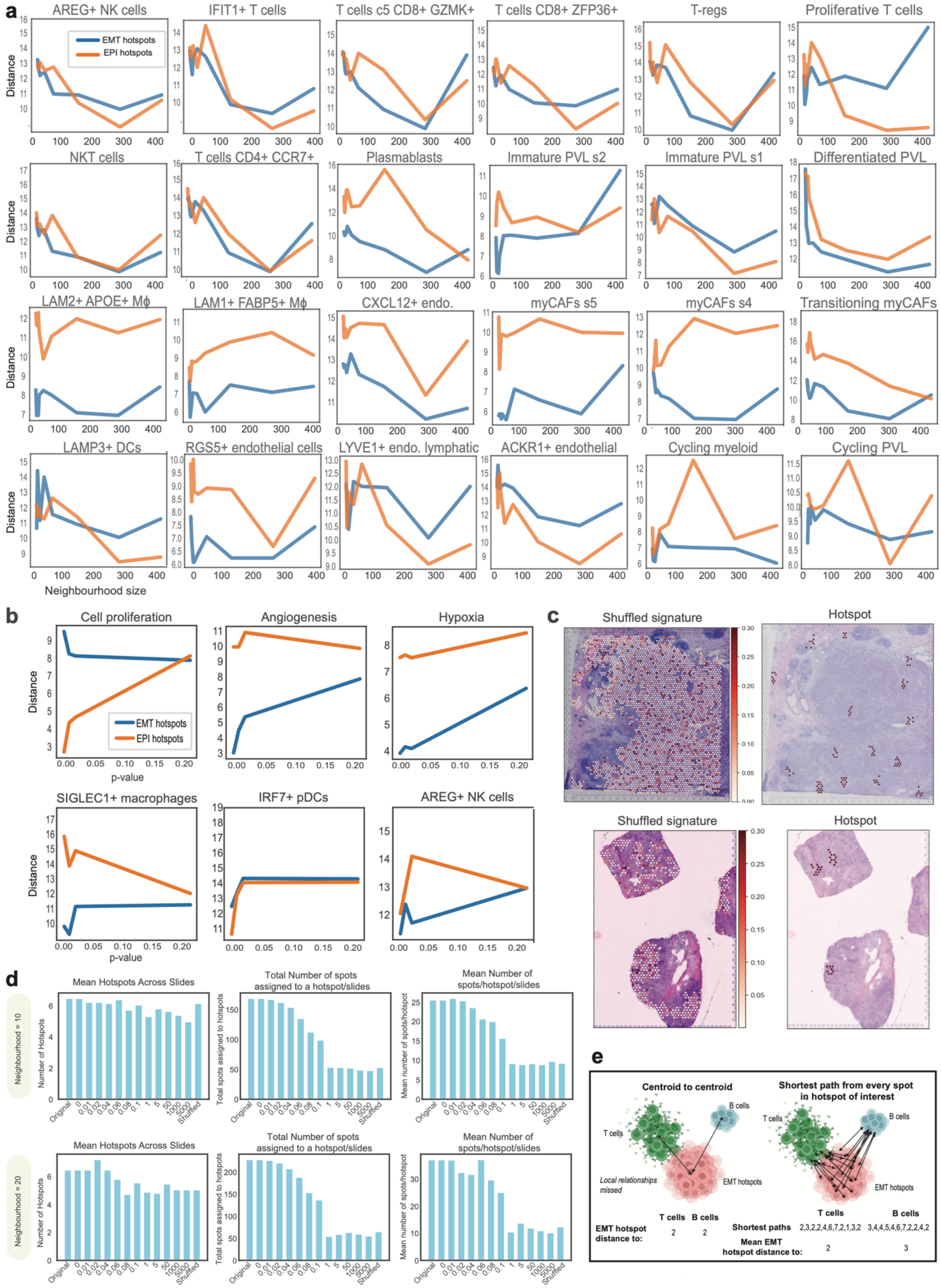


**Supplementary Figure 3: Hotspot sensitivity analysis. (a)** Sensitivity plots highlighting the distance from EMT hotspots (blue) and EPI hotspots (orange) to various TME components as the hotspot size increases. Distances are averaged over all 12 slides. (**b**)

Sensitivity plots highlighting the distance from EMT hotspots (blue) and EPI hotspots (orange) to various TME components as the p-value parameter using to detect statistically significant hotspots increases. Distances are averaged over all 12 slides. **(c)** Spatial plots of shuffled EMT signature (left) and hotspots produced from shuffled signature (right) for Slide 6 (top) and Slide 7 (bottom). **(d)** Frequency bar plots for neighbourhood size=10 hotspots (top) and neighbourhood size=20 hotspots (bottom), highlighting the average number of hotspots across a slide (left), total number of spots assigned to a hotspot/slide (middle) and average number of spots/hotspot/slide (right) with increased amounts of noise added to the EMT signature, or spot shuffling. **(e)** Diagram highlighting key differences between the centroid distance approach and the shortest path distance approach. The shortest path approach captures more local variation and therefore detects a difference between the T-cell and B-cell hotspots shown in this example (distances shown for illustration purposes). The centroid-to-centroid approach however would be unable to capture this.

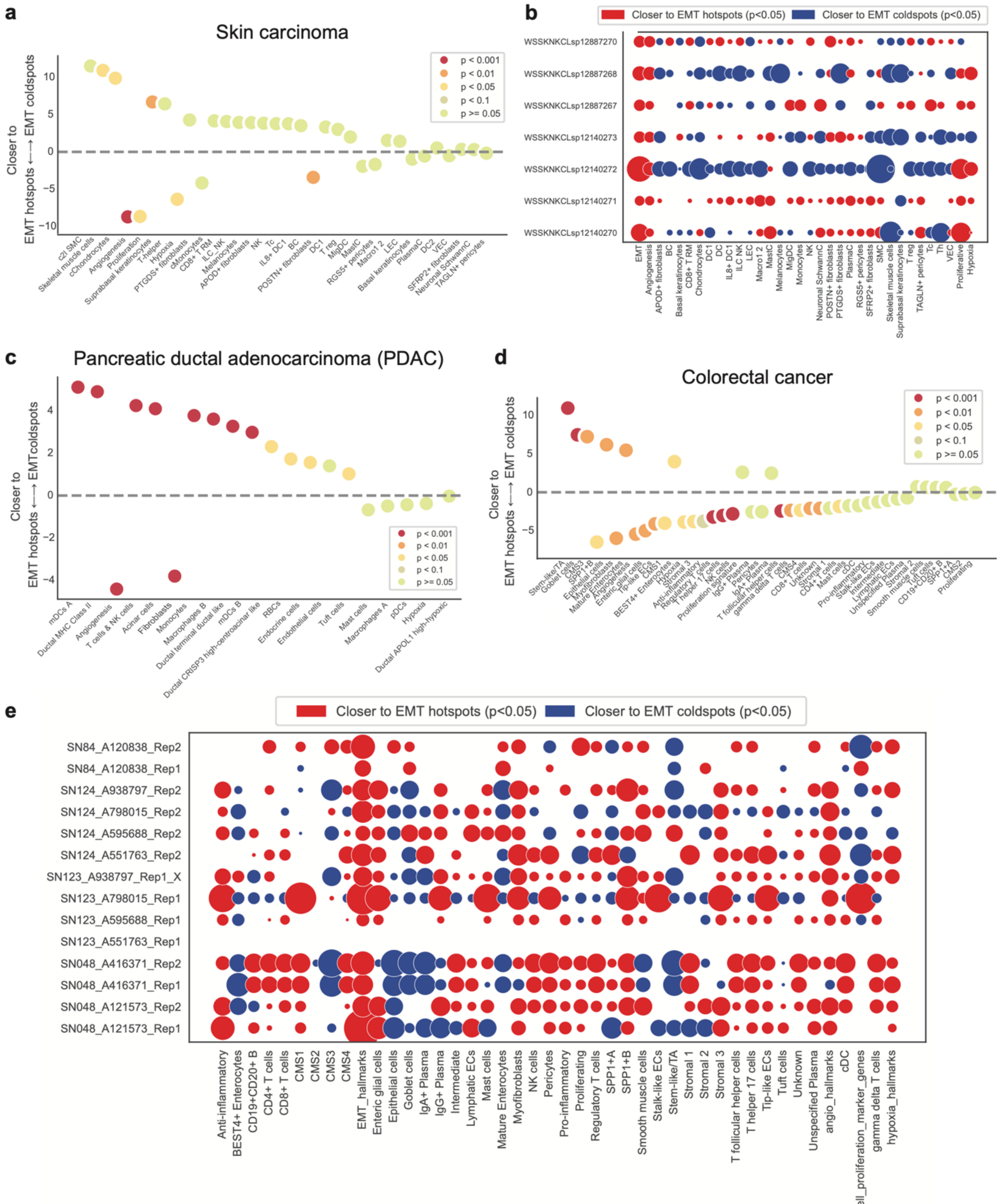


**Supplementary Figure 4: EMT hotspot analysis in other cancer types. (a)** Distances from various cells in the TME to EMT hot/cold regions in basal cell skin carcinoma. The dashed line represents no difference in proximity to either EMT hotspots or EMT coldspots. The dots situated to the right of the dashed line indicate cell populations that are significantly closer to EMT hotspots, ordered by decreasing proximity. The colours indicate the p-value ranges obtained from the GEE model fit. **(b)** Bubble plot depicting distances between cancer hallmark signatures and TME classes and EMT hotspots/coldspots for each BCC slide (row). Blue depicts hallmarks that are significantly closer to EMT coldspots and red represents hallmarks that are significantly closer to EMT hotspots (Student's t test $p<0.05$), adjusted for multiple testing using the Bonferroni correction. White indicates a non-significant relationship. **(c)** Similar to (a) for one PDAC sample. **(d)** Similar to (a) for colorectal cancer slides. **(e)** Similar to (b) for the colorectal cancer slides.

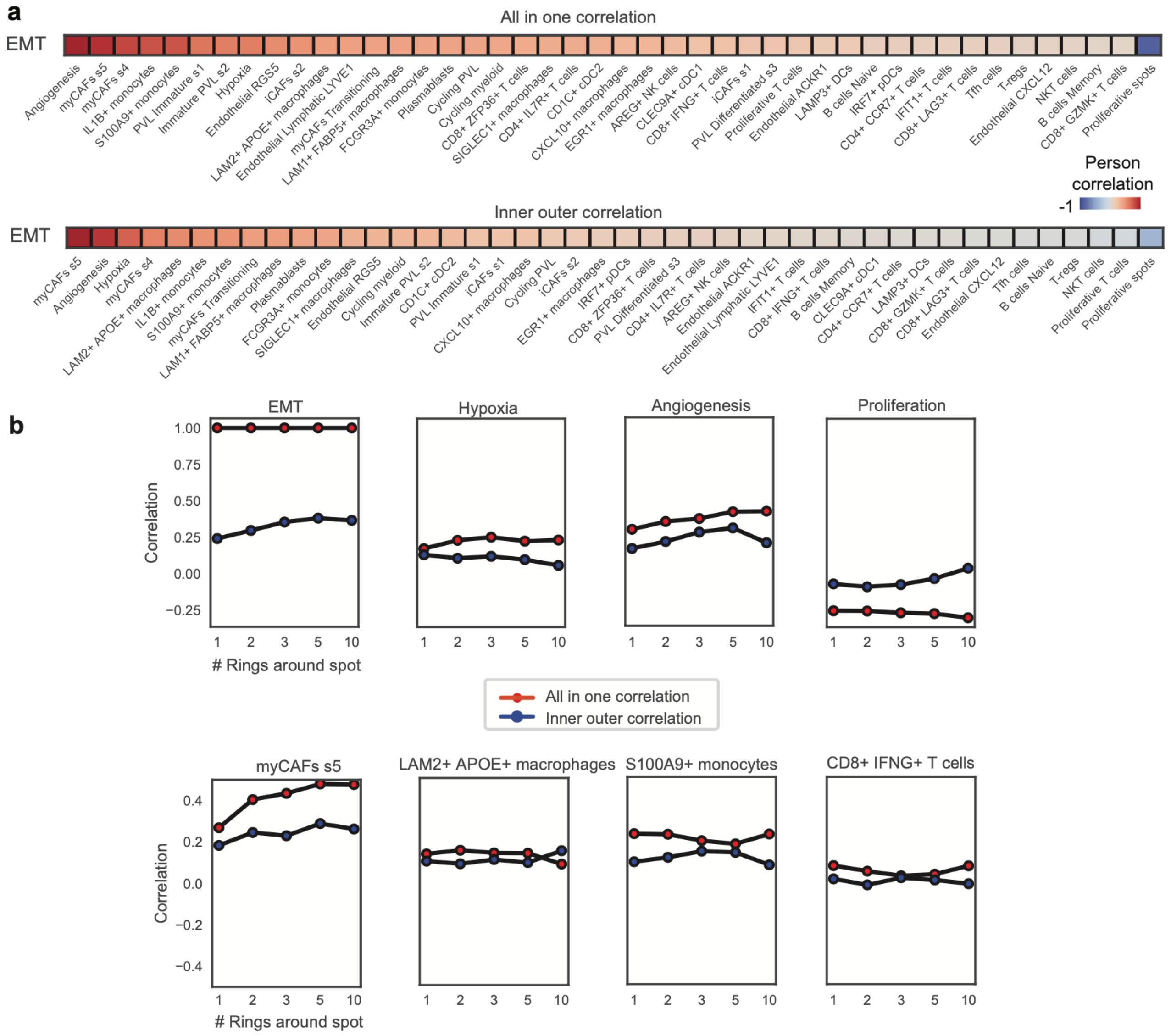


**Supplementary Figure 5: Neighbourhood enrichment analysis of EMT spatial dynamics. (a)** Neighbourhood enrichment analysis results employing all in one correlation (top) and inner outer correlation (bottom) approaches. The squares display correlations between EMT levels within tumour cell spots and the abundance of various cell populations within the immediate TME (surrounding spots only). Red indicates a positive correlation, blue a negative correlation and white a non-significant correlation (Pearson p>0.05). 1 ring is used to define the neighbourhood. **** p<0.0001, *** p<0.001, ** p<0.01, * p<0.05. **(b)** Line plots illustrating the impact of progressively expanding the number of concentric rings - from 1 to 10 - around a Visium spot on the correlation between the EMT signature and various cells in the TME. Each ring represents an incremental distance from the central spot and encompasses the surrounding spatial transcriptomic spots.

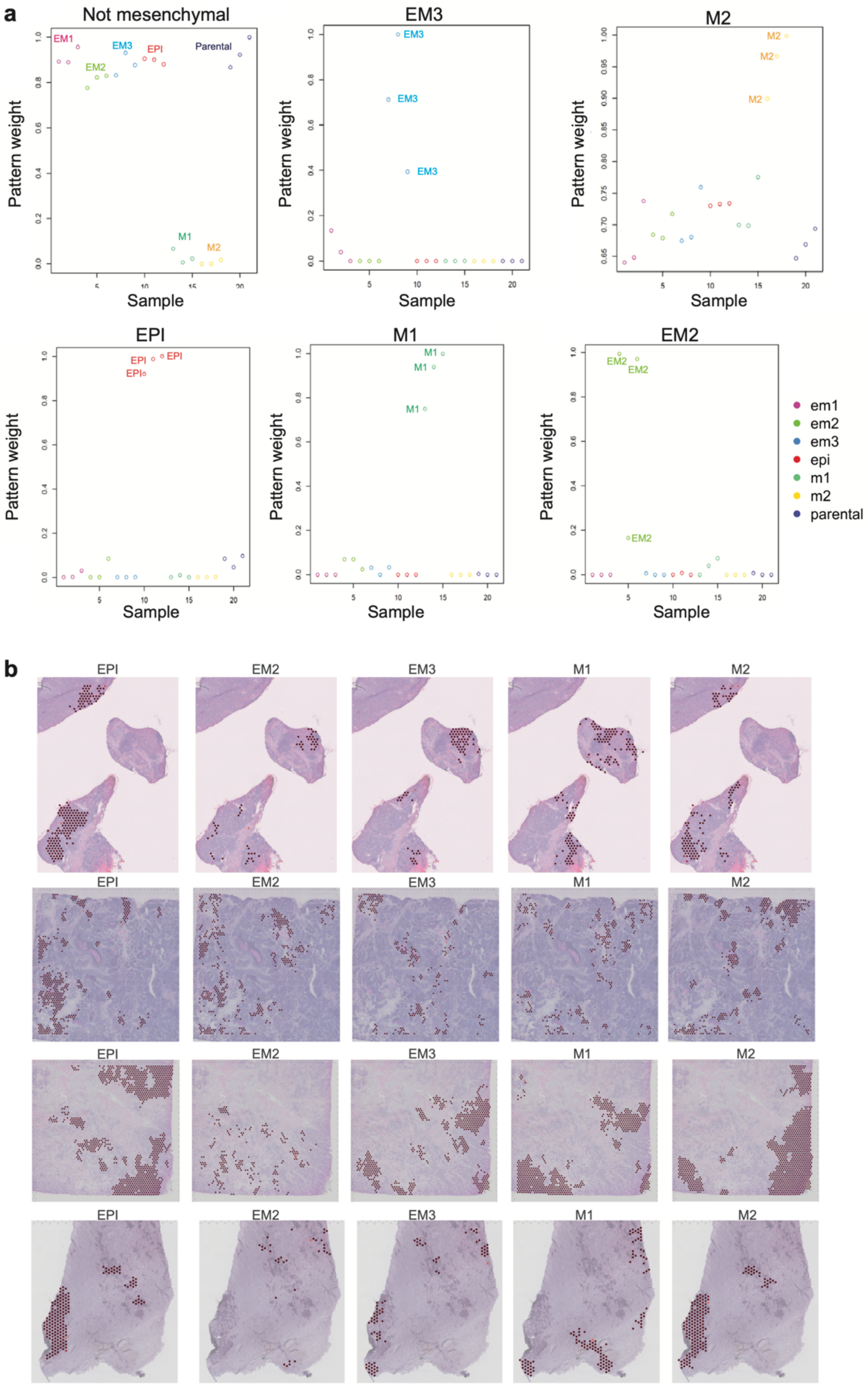


**Supplementary Figure 6: EMT state capture in spatial transcriptomics slides. (a)** NMF patterns captured from Brown et al[25], consisting of seven RNA-seq sequenced cell clones, with three repeats spanning the EMT spectrum including epithelial-like (EPI), quasi-mesenchymal (M1), fully mesenchymal (M2) and three distinct intermediates (EM1, EM2, EM3). Each circle corresponds to one cell clone from the original dataset, and is coloured according to the assigned state. The pattern weights for each cell clone are plotted for each pattern. The patterns that were able to separate the cell clones are annotated. **(b)** Spatial plots highlighting the distribution of EMT state hotspots (EPI, EM2, EM3, M1 and M2). Every row corresponds to one Visium slide.

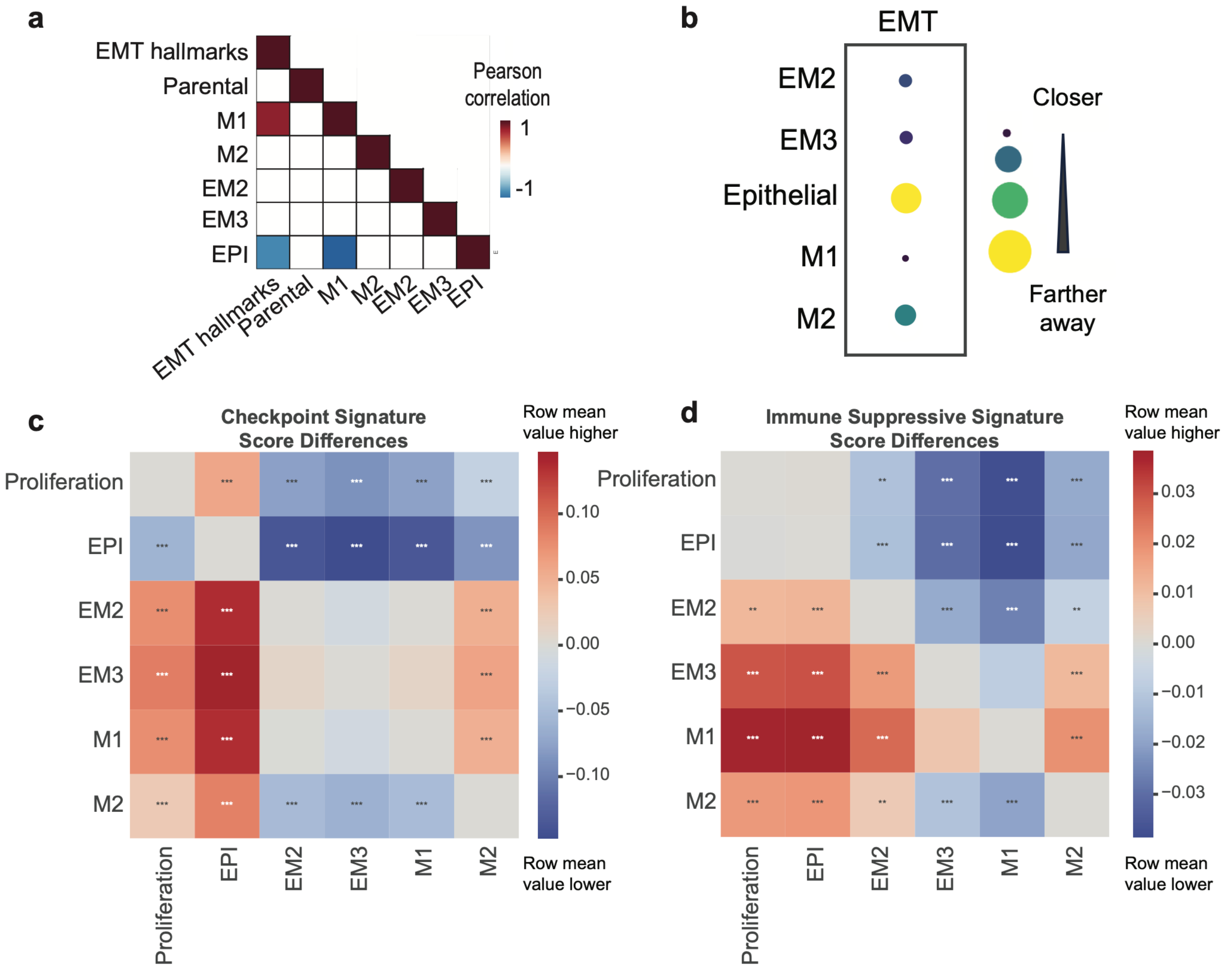


**Supplementary Figure 7: EMT state correlations within spatial transcriptomics slides.** **(a)** Correlation plot of EMT state scores across all spots and slides. Red indicates positive correlation; blue indicates negative correlation. Only significant correlations (p<0.05) are shown. White squares indicate non-significant correlations. **(b)** Bubble plot depicting the mean distance from individual EMT state hotspots to the EMT hallmark hotspots defined in our original analysis. Smaller bubbles represent shorter distances. **(c)** Heatmap representing differences in immune suppression signature scores for EMT state hotspots. Red indicates that the EMT state in the row has a higher signature score than the EMT state in the column, blue indicates that the EMT state in the row has a lower score. Stars depict the significance of the difference (Student's t test Bonferroni adjusted p-values are shown, *** p<0.001, ** p<0.01, 8 p<0.05). **(d)** Heatmap representing differences in checkpoint signature scores for EMT state hotspots. Red indicates that the EMT state in the row has a higher mean score for this signature than the EMT state in the column, blue indicates that the EMT state in the row has a lower mean score. Stars depict the significance of the difference (Student's t test Bonferroni adjusted p-values are shown, *** p<0.001, ** p<0.01, * p<0.05).
